# Scrutinization on Docking Against Individually Generated Target Pockets for Each Ligand

**DOI:** 10.1101/2025.01.01.630989

**Authors:** Rui Zhan, Wenyi Zhang, Jing Huang

**Affiliations:** Key Laboratory of Structural Biology of Zhejiang Province, School of Life Sciences, Westlake University, Hangzhou, 310024, China; Westlake AI Therapeutics Lab, Westlake Laboratory of Life Sciences and Biomedicine, Hangzhou, 310024, China; Institute of Biology, Westlake Institute for Advanced Study, Hangzhou, 310024, China

**Keywords:** Virtual Screening, Structure Prediction, Ensemble Docking, Individual Docking

## Abstract

The longstanding challenge of limited exploration in protein receptor conformational space continues to constrain the precision of molecular docking. Ensemble docking, which employs methods such as molecular dynamics simulations to generate multiple receptor conformations for docking, has improved accuracy but remains limited by incomplete sampling and an inability to fully account for ligand-induced fit. To address these limitations, we introduce the concept of individual docking, a novel approach that involves docking against receptor conformations generated individually for each ligand in the docking library. This approach has only very recently become feasible due to advances in protein structure prediction, in particular end-to-end protein-ligand complex prediction technologies. In this study, we performed individual docking on 27 targets from the DUD-E dataset, using a two-step protocol that integrates NeuralPLexer or AlphaFold3 for receptor conformation sampling, followed by physics-based docking. Our results reveal that individual docking with AlphaFold3 predictions yields more than a twofold improvement in enrichment factors compared to standard docking. Furthermore, individual docking recovers distinct sets of active ligands, thereby expanding the diversity of virtual screening hits. Detailed analyses of pocket and ligand conformations suggest several potential incompatibilities between deep learning-based and physics-based virtual screening tools.

## Introduction

High-throughput screening has long been the standard method for identifying biologically active compounds, but its application is often limited by high costs. Virtual screening (VS), which simulates compound-target interactions or evaluates compound and pharmacophore similarities computationally, has emerged as a powerful alternative [1–3]. VS has been widely adopted in drug discovery, not only for accelerating early-stage drug discovery but also for applications such as drug repurposing [4], compound optimization [5], and target identification [6]. With the advancement of protein 3D structure determination methods, structure-based virtual screening, such as molecular docking and free energy perturbation calculations, has gained increasing attention [7].

Molecular docking computationally positions molecules within a selected binding pocket through search algorithms, generating binding poses by considering factors such as pocket properties, ligand rotations, translations, and internal degrees of freedom. By employing empirical scoring functions, it further provides a docking score to estimate the complementarity between ligands and targets, enabling the effective ranking of millions of small molecules in virtual libraries. Docking is probably the most popular and successful computational approaches in drug discovery [8]. Early research on molecular docking focused on improving scoring functions and ligand sampling methods. Advancements in scoring functions have sought to enhance docking accuracy by incorporating key structural and physicochemical properties [9], while improvements in sampling techniques have involved iterative random search algorithms such as Monte Carlo and evolutionary algorithms [10,11]. However, a remaining limitation of docking is that the protein structure is kept fixed, which does not account for protein dynamics. Even in flexible docking, only a limited number of side chains can be rotated.

One way to partially account for protein flexibility is ensemble docking [12–14], which involves performing docking calculations against multiple receptor conformations simultaneously. To generate a diverse and representative set of receptor conformations, molecular dynamics (MD) simulations or normal mode analysis of the apo-protein are often employed. While the conformational change induced by ligand binding is often described using the induced fit model, the conformational selection model can also be considered, which suggests that a protein populates all binding-necessary conformations even in its unbound state [15–17]. However, obtaining a convergent protein conformational landscape remains highly challenging even with enhanced sampling techniques [18]. Moreover, selecting a limited number of conformations from the landscape to perform ensemble docking for a given screening library is equally difficult.

Artificial intelligence (AI), in particular deep learning (DL) methods, offer exciting new approaches to addressing challenges in structure-based drug design such as the scarcity of high-quality protein structures and the limited exploration of protein conformational space [19–24]. Most notably, AlphaFold2 enables highly accurate end-to-end protein structure prediction using its Evoformer module within the transformer architecture, combined with geometric modeling. The AlphaFold [25], AlphaFold2 [26], and contemporary methods [27–32] have primarily focused on predicting protein-only structures, albeit they were trained on PBD entries that often correspond to ligand-bound conformations. In the 2022 Critical Assessment of Structure Prediction (CASP15) competition [33], protein-ligand complex structure prediction was introduced for the first time. The winning method, CoDock [34], combined template-based docking on AlphaFold2-predicted apo structures with deep learning-based scoring to assess ligand binding predictions.

Diffusion models, inspired by thermodynamics modeling, represent the current state-of-the-art in DL generative models. By learning an estimated score function, diffusion models effectively reduce the dependence on large training datasets. Diffusion models further advance structure prediction by enabling more accurate predictions of biomolecular complexes, including protein interactions with small molecules, metals, nucleic acids, and other biomolecules [35–37]. A wide range of diffusion-based generative models has emerged for predicting the structure of protein-ligand complexes, including AlphaFold3 [38], RoseTTAFold AA [39], NeuralPLexer [40,41], Chai-1 [42], and Boltz-1 [43]. Unlike physics-based docking methods, these DL models perform blind predictions without requiring prior knowledge of the binding pocket and generate fully induced-fit conformations of the target proteins. However, DL models fail to differentiate between binders and non-binders, as they always predict complex structures based on given inputs and lack the capability to directly predict binding affinities. The integration of structure prediction, docking, and virtual screening within a unified computational or DL framework has emerged as a central focus in advancing ligand-protein interaction prediction [44,45]. To this end, effective scientific validation is critical for assessing and understanding the compatibility between physics-based and DL methods, ensuring their synergistic integration.

In this study, we propose an individual docking (ID) pipeline that leverages a DL model to predict protein-ligand complex structures and then performs molecular docking using the predicted protein conformations, which can be regarded as individually generated pocket for each ligand. To evaluate the performance of this approach, we conducted virtual screening on 27 systems from the DUD-E (a database of useful decoys: enhanced) dataset [46], encompassing a total of 891,460 (445,730 for each DL model) predicted protein-ligand complex structures generated by NeuralPLexer (NP) and AlphaFold3 (AF3) model. To assess the flexibility of the binding pockets, we used a 3D pocket structure alignment tool to evaluate the generated conformations. In addition, we analyzed the docking score distribution and compared the predicted ligand poses from DL prediction with the post-docking poses. Through this comprehensive evaluation, we provide insights into the compatibility between DL-based structure prediction and physics-based docking methods, highlighting the potential advantages and limitations of integrating DL models with conventional molecular docking techniques for virtual screening.

## Results

### Docking calculations have to be performed on individually generated pockets by NeuralPLexer

The 27 systems from the DUD-E dataset were used to investigate ligand-protein interactions with individually generated pockets for each ligand, as used in a previous study benchmarking AlphaFold2-predicted structures for hit discovery [47]. For individual docking, we employed NeuralPLexer to predict ligand-protein complex structures, taking the protein sequence and ligand graph structures as inputs. Subsequently, one can directly compute the docking score using an empirical scoring function, such as Vina [48], to evaluate the interaction based on the ligand-protein conformation predicted by NeuralPLexer. Despite retaining most of the information from the structure prediction, direct scoring performed poorly, with most Vina scores being positive, indicating unreasonable binding as assessed by the Vina scoring function. Among 445,730 cases, 96.42% of the docking scores exceeded 0 kcal/mol, and 10.89% were greater than 100 kcal/mol (Fig. S1A). Notably, only 1275 complex structures directly generated by NeuralPLexer achieved docking scores within the typically accepted range of below -6 kcal/mol, accounting for a mere 0.29% (Fig. S1B). Direct calculation of the docking scores suggests a lack of stringent physical constraints in the predicted complex structures, such that some atomistic distances between the protein and ligand can be excessively close and may lead to steric clashes. Consequently, docking calculations, which involve sampling ligand binding poses in the pocket, must be performed to obtain valid docking conformations for evaluating ligand-protein interactions using docking scores.

To further investigate the impact of NeuralPLexer sampling on virtual screening, we designed two groups of individual docking experiments with progressively decreasing levels of information retention from NeuralPLexer: individual rigid docking (ID-rigid-NP) and individual docking (ID-NP). In ID-rigid-NP, Vina was used to resample the ligand docking orientations while both the protein and ligand conformations were fixed. In contrast, ID-NP preserved only the receptor conformation and allowed the ligand conformations and docking poses to be resampled during the docking calculations. The binding site was determined based on the ligand’s predicted position from NeuralPLexer. Standard docking (SD) was performed as the baseline using the crystal structures of the receptors. Virtual screening (VS) performance of individual docking was benchmarked and compared with SD across 27 systems from the DUD-E dataset.

### Comparison of VS performance by NeuralPLexer

The enrichment factors (EFs) at 1% cutoff are shown in Table 1, comparing ID-NP with SD. In nine out of 27 systems studied, for example ACES and AKT2, individual docking led to increased enrichment in active hits when selecting the top 1% of compounds. In another 16 systems, ID-NP showed worse performance compared with SD, while in two systems (AMPC and HSP90A) neither ID-NP nor SD identified any active hits at the 1% cutoff. The median enrichment factors for active compounds at the top 1%, 5%, and 10% ranges were 2.35, 2.16, and 2.05 for ID-NP, slightly but consistently lower than the corresponding values for SD, which were 3.08, 2.77, and 2.65, respectively (data for the 5% and 10% cutoff are provided in Table S1).

**Table 1.**
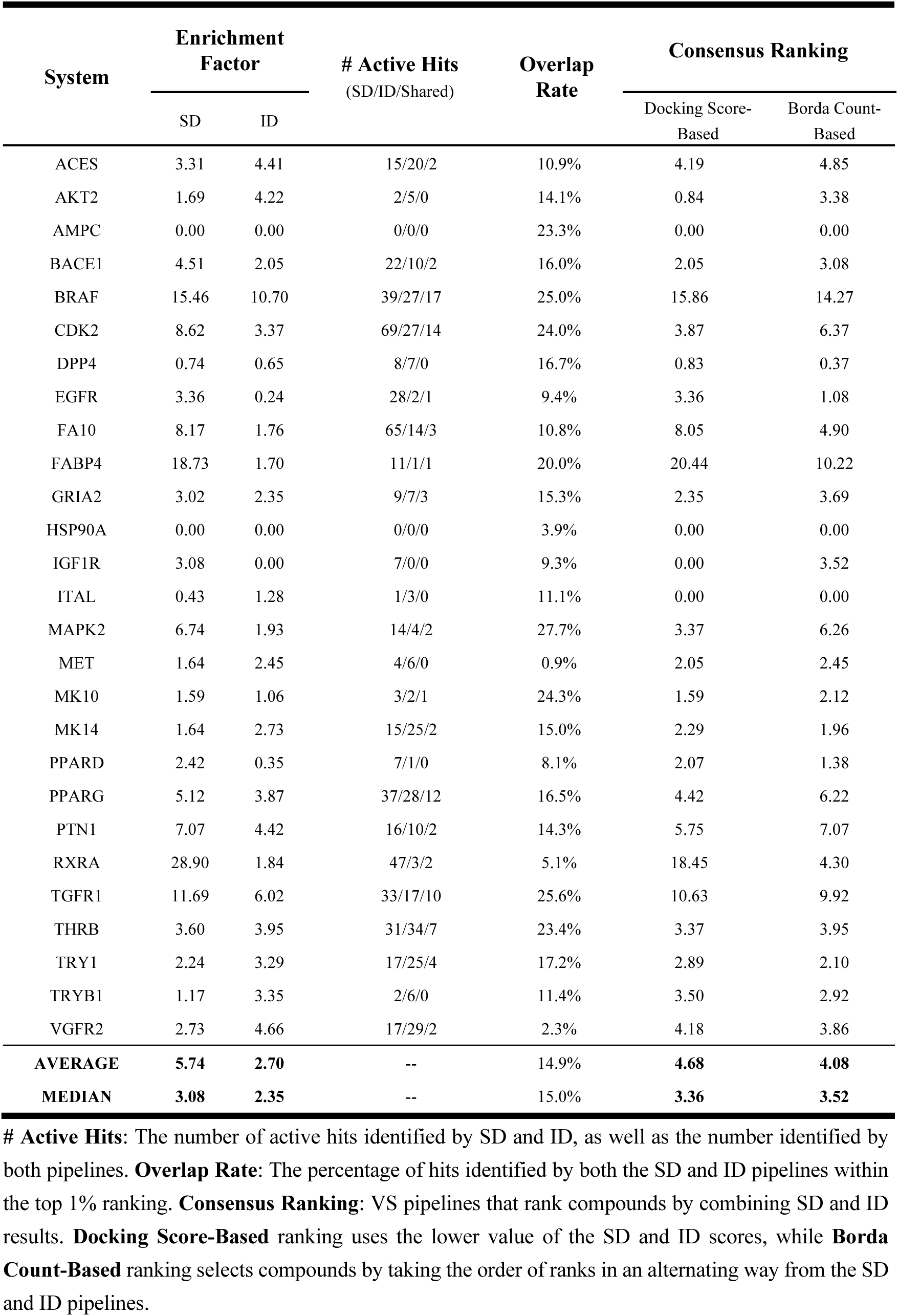
Virtual screening performance of ID with NP evaluated at 1% cutoff.

In addition to the EFs, we also calculated the area under the ROC curve (AUC) values (Fig. 1 and S2). The average AUC value for ID-NP across all 27 systems studied was 0.62, which was 0.05 lower than the AUC value of 0.67 for SD. The ROC curves aligned with the 1% EF results, showing that SD generally had better VS performance. However, in a few systems, such as TRY1 and VGFR2, ID-NP demonstrated a stronger capability to select active hits. We also note that while in some systems (e.g., MAPK2 and THRB) the ID-NP and ID-rigid-NP pipelines produced almost identical results, in most cases, their performance differed significantly.

**Figure 1.**
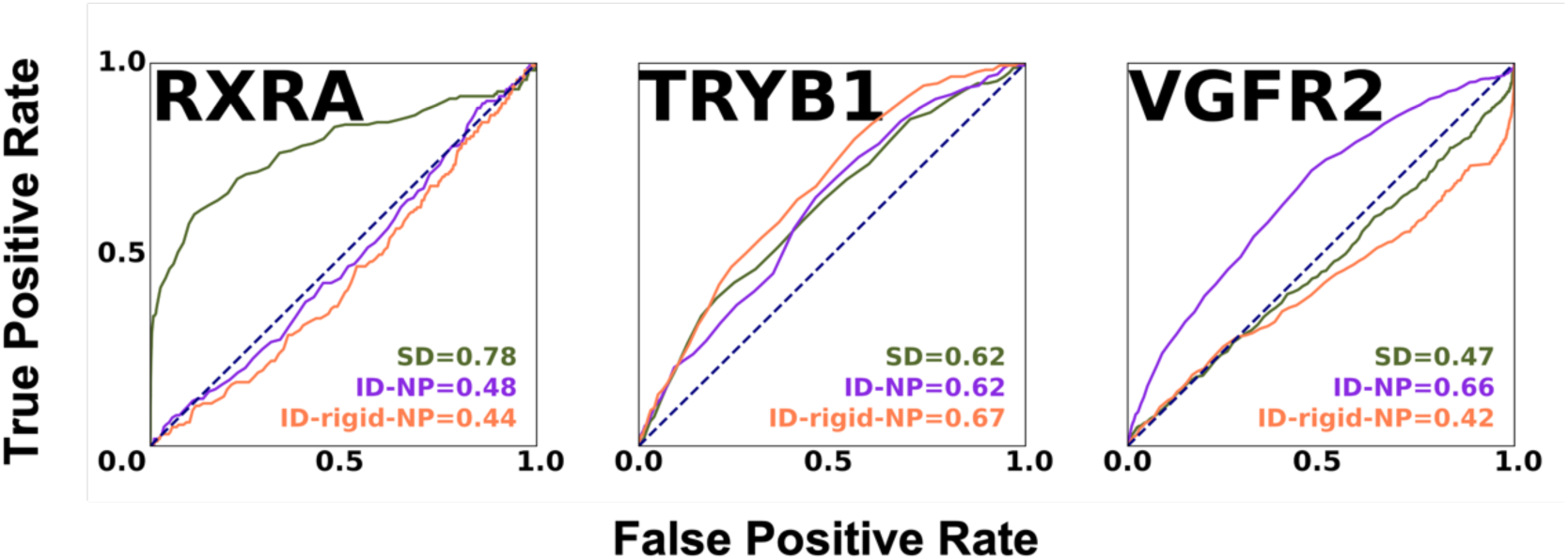
ROC curves and AUC values for RXRA, VGFR2, and TRYB1 systems, which are representative for standard, individual, and individual-rigid docking with NeuralPLexer pipelines, respectively.

We also examined the active hits identified from the top 1% ranked compounds using ID-NP and SD and found significant differences in the compounds predicted by the two pipelines (Table 1). For example, in the ACES system, SD identified 15 active hits and ID-NP identified 20, with only two compounds overlapping. In DPP4, SD identified eight active hits and ID-NP found seven, but none of these 15 hits overlapped. The BRAF system had the highest number of shared active hits, with SD identifying 39 active hits and ID-NP detecting 27, 17 of which were shared. Notably, this still means that individual docking identified 10 unique active hits compared to standard docking. This echoes a recent observation that docking calculations using AlphaFold2-predicted target conformations achieve similar VS performance compared to using experimentally determined structures, but the identified active hits are markedly different [49]. Taking decoys into account, the overlap between the top 1% SD and the top 1% ID-NP compounds ranged from 0.9% to 27.7%, with a median of 15.0%. Such a low overlap rate indicates that, even with identical docking settings, using either a single experimental target conformation or a library of individually generated conformations results in the prioritization of very different compounds. The origin of this discrepancy will be analyzed in the following subsections.

### Pocket similarity and flexibility sampled by NeuralPLexer

We evaluated the differences between the predicted and crystal pockets using the pocket shape alignment tool called PPS (Quick & accurate Protein Pocket Structural Alignment) [50]. The PPS score, ranging from 0 to 1, reflects the similarity between the predicted and crystal pockets, with a score closer to 1 indicating a high resemblance and a score closer to 0 suggesting greater pocket flexibility. In total, we used NeuralPLexer to predict ligand-protein complexes for 27 systems from the DUD-E dataset, generating 445,730 conformation samples. The computed pocket similarity scores (PPS scores) and root mean square deviations (RMSDs) for each sample are shown in Fig. 2. The average and median PPS scores were 0.59 and 0.61, respectively, while the average and median RMSD values were 1.08 Å and 1.06 Å, respectively. Of these, 24,219 pockets had PPS scores below 0.4 and RMSD values greater than 1.5 Å. Notably, the majority of these pockets were identified in the DPP4 system, which contributed 17,428 cases, followed by VGFR2, with 1,932 samples.

**Figure 2.**
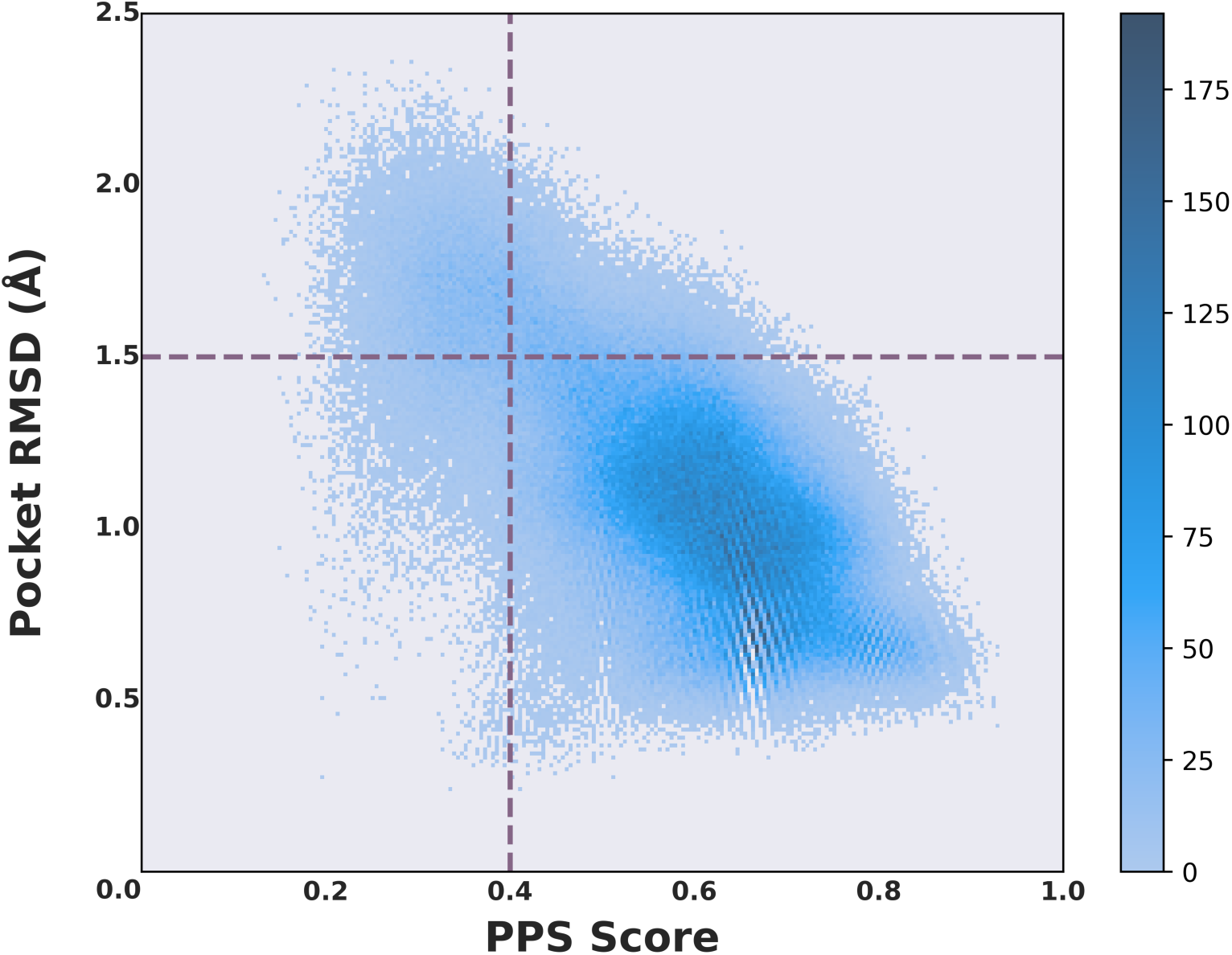
Distribution of PPS scores and pocket RMSD values for pockets generated by NeuralPLexer. The PPS scores were calculated using the PPS-align method, which aligns the generated pockets with the corresponding crystal pocket.

Interestingly, for these systems, the enrichment factors increased with individual docking, either directly (e.g., VGFR2, which increased from SD 2.73 to ID-NP 4.66) or indirectly (e.g., DPP4, where SD and ID-NP identified completely different active hits such that consensus ranking could be beneficial; see below). Most of the remaining systems produced approximately 10-150 conformation samples with similar characteristics, suggesting relatively low levels of pocket flexibility. We also calculated the Pearson’s correlation coefficient (PCC) between the PPS score and the number of heavy atoms in the ligand. The PCC value was 0.08, indicating that the dynamic pocket generation in NeuralPLexer was not influenced by ligand size. This result highlights that the DL-based model can effectively handle pocket flexibility, adapting to variations in pocket geometry regardless of ligand size.

### Comparison of docking scores

We further analyzed the docking scores obtained with SD, ID-NP and ID-rigid-NP, which include approximately 440,000 docking cases for each pipeline. Significant differences in the distribution of docking scores were observed, as shown in the histogram plot in Fig. 3A. Interestingly, ID-rigid-NP resulted in a significantly larger number of cases with very favorable binding (docking scores below -11 kcal/mol, Fig. 3B), as well as a greater number of cases with very unfavorable binding (positive docking scores). Essentially, ID-rigid-NP adjusts the predicted binding pose through rigid-body translation and rotation of the ligand, while ID-NP further allows for ligand deformation and internal rotation along rotatable bonds. A large number of positive docking scores associated with ID-rigid-NP indicates cases where the docking program was unable to identify a reasonable binding pose given the conformation of the protein receptor. It’s worth noting that, in contrast, ID-NP resulted in much fewer unfavorable binding cases compared to SD, although in general ID-NP and SD had similar docking score distributions, with ID-NP showing a slightly higher probability in the [-14, -12] and [-7, -4] kcal/mol ranges.

**Figure 3.**
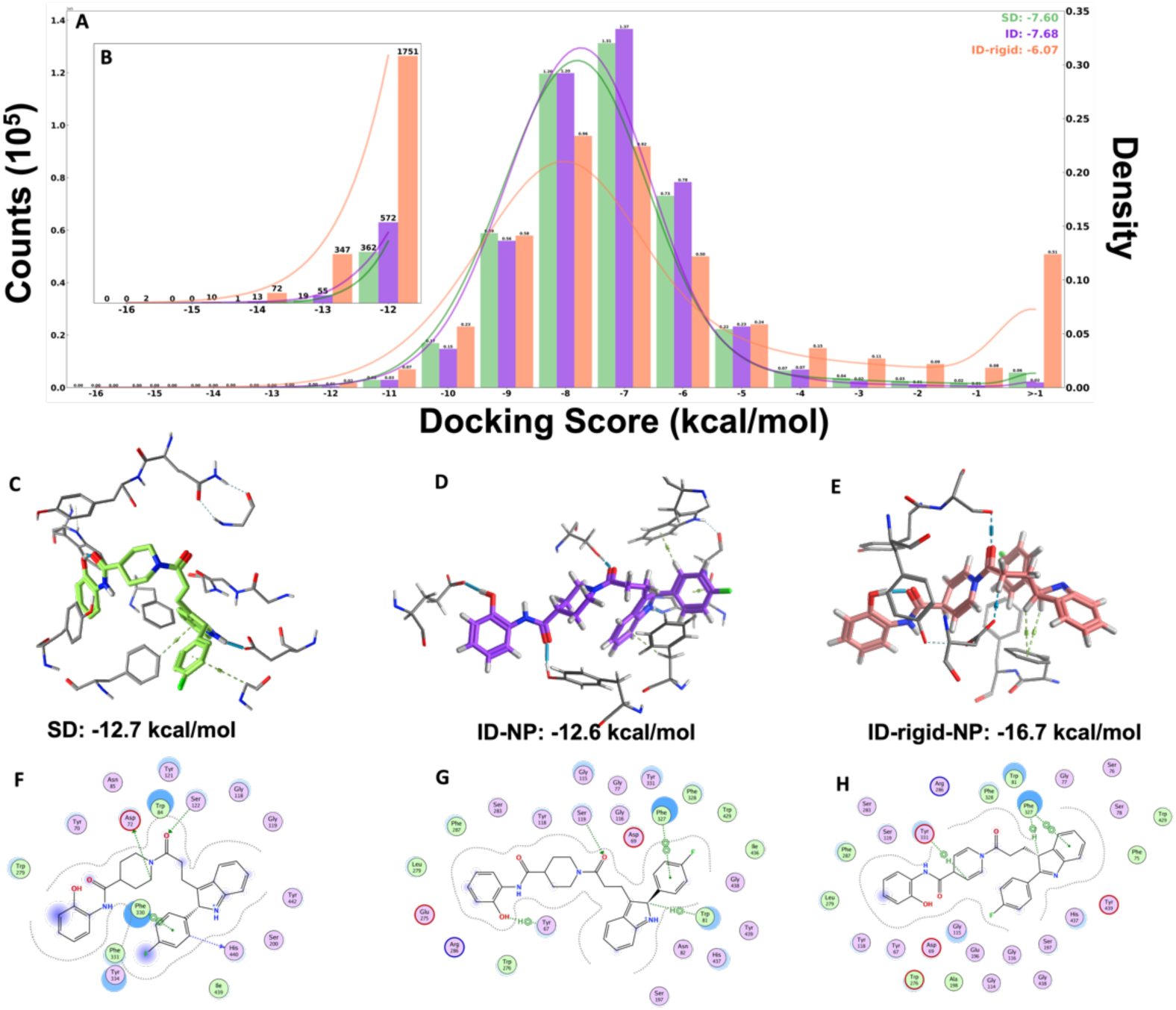
Comparison of docking score distributions for the SD, ID-NP, and ID-rigid-NP pipelines. A: Overall docking score distribution for SD, ID-NP, and ID-rigid-NP. B: Docking score distribution for scores below -11 kcal/mol. C-E: Docking poses and interactions of ZINC63182445 targeting ACES, obtained with SD, ID-NP, and ID-rigid-NP, respectively. F-H: Detailed interaction diagrams for SD, ID-NP, and ID-rigid-NP, respectively.

A zoom-in on the favorable binding range is shown in Fig. 3B. ID-rigid-NP computed 9,167 cases with docking scores below -11 kcal/mol, approximately 2.6 times more than SD (3,345 cases) and ID-NP (3,591 cases). Some ID-rigid-NP docking scores were lower than -15 kcal/mol, while no such cases were observed with SD or ID-NP, suggesting that the DL model’s sampling process could optimize docking scores for certain ligands. Specifically, we selected the case with the most favorable docking score with ID-rigid-NP for conformational analysis, as shown in Fig. 3C-H. This case involves the docking result of the 22,573rd decoy ligand (ZINC63182445) targeting ACES. The docking scores were -12.7 kcal/mol for SD, -12.6 kcal/mol for ID-NP, and -16.7 kcal/mol for ID-rigid-NP.

By comparing the docking poses of SD and ID-rigid-NP, we found that while SD identified four hydrogen bond interactions, the ligand underwent significant bending to fit the pocket shape, resulting in a high-energy conformation and consequently poorer docking scores. In contrast, the ID-rigid-NP docking pose demonstrated a significant adjustment in pocket shape due to NeuralPLexer sampling (PPS score: 0.64, pocket RMSD: 1.15 Å), creating a more compatible fit with the ligand conformation. This adjustment facilitated an aromatic interaction network with the ligand through the critical residue F330. Despite having fewer hydrogen bond interactions than SD, ID-rigid-NP achieved a substantial improvement in docking score, showcasing the advantage of DL-sampled protein pocket conformations in improving docking performance. Moreover, docking into the same pocket resulted in overall similar ligand-protein binding modes between ID-NP and ID-rigid-NP. However, a twist at one end of the ligand, with the fluorobenzene moiety making contact with F330 instead of the indole moiety, effectively reset the docking score to -12.6 kcal/mol (Fig. 3, G and H). This example demonstrates the sensitivity of docking scores to subtle changes in ligand orientation and interactions within the given pocket.

To further investigate the significance of more favorable docking scores in VS performance, we analyzed the proportion of active ligands in a subset of 9,167 cases identified by ID-rigid-NP with docking scores below -11 kcal/mol. Among these, only 309 were active ligands, accounting for a mere 3.4%, which corresponds to an enrichment factor of 1.31. This is notably lower than the 1% EFs achieved by any of the three pipelines. Therefore, the improvement in docking scores associated with individual rigid docking does not translate into enhanced virtual screening performance. We also compared the docking poses and NeuralPLexer-predicted poses of compounds in this subset using the DockRMSD method. DockRMSD is a convenient tool for comparing ligand structures without requiring RMS alignment of the target conformation, allowing efficient calculation of the RMSD values between two ligand sets [51]. Relatively large DockRMSD values, with an average of 5.81 Å and a median of 7.74 Å, were observed. This indicates that, despite preserving receptor and ligand conformations from DL predictions in ID-rigid-NP, most cases with improved docking scores still relied on docking algorithms to search for significantly different ligand orientations.

We also performed a direct comparison of docking scores for each ligand between SD and ID-NP to explore whether the docking scoring can be changed by introducing DL sampling, as illustrated by the scatter plots in Fig. 4. Correlation coefficients were calculated. Only docking scores below -4 kcal/mol were considered. For most systems, docking scores obtained using two different types of target conformations correlated well, with correlation coefficients ranging from 0.15 (VGFR2) to 0.87 (AMPC). The median correlation coefficient was 0.57. Four systems (PTN1, RXRA, MET, and VGFR2) exhibited particularly low R values, suggesting more substantial influences of DL sampling on docking scores. However, it remains unclear these shifts lead to improvements or declines in VS performance: SD outperformed ID-NP in PTN1 and RXRA, while the reverse was observed in MET and VGFR2. Furthermore, among the 427,094 cases included in the analysis, 210,129 (49.2%) had docking scores that favored ID-NP. This observation aligns with our findings in Fig. 3A, where the two methods exhibited comparable docking score distributions. Furthermore, among the four systems where ID-NP scores were consistently more negative than SD scores (BACE1, CDK2, GRIA2 and MK14), only MK14 demonstrated an improvement in EF with the ID-NP pipeline. This is consistent with our analysis of the ID-rigid-NP favorable subset, where more favorable docking scores did not correlate with an enhancement in virtual screening capability, which requires differentiating actives from decoys.

**Figure 4.**
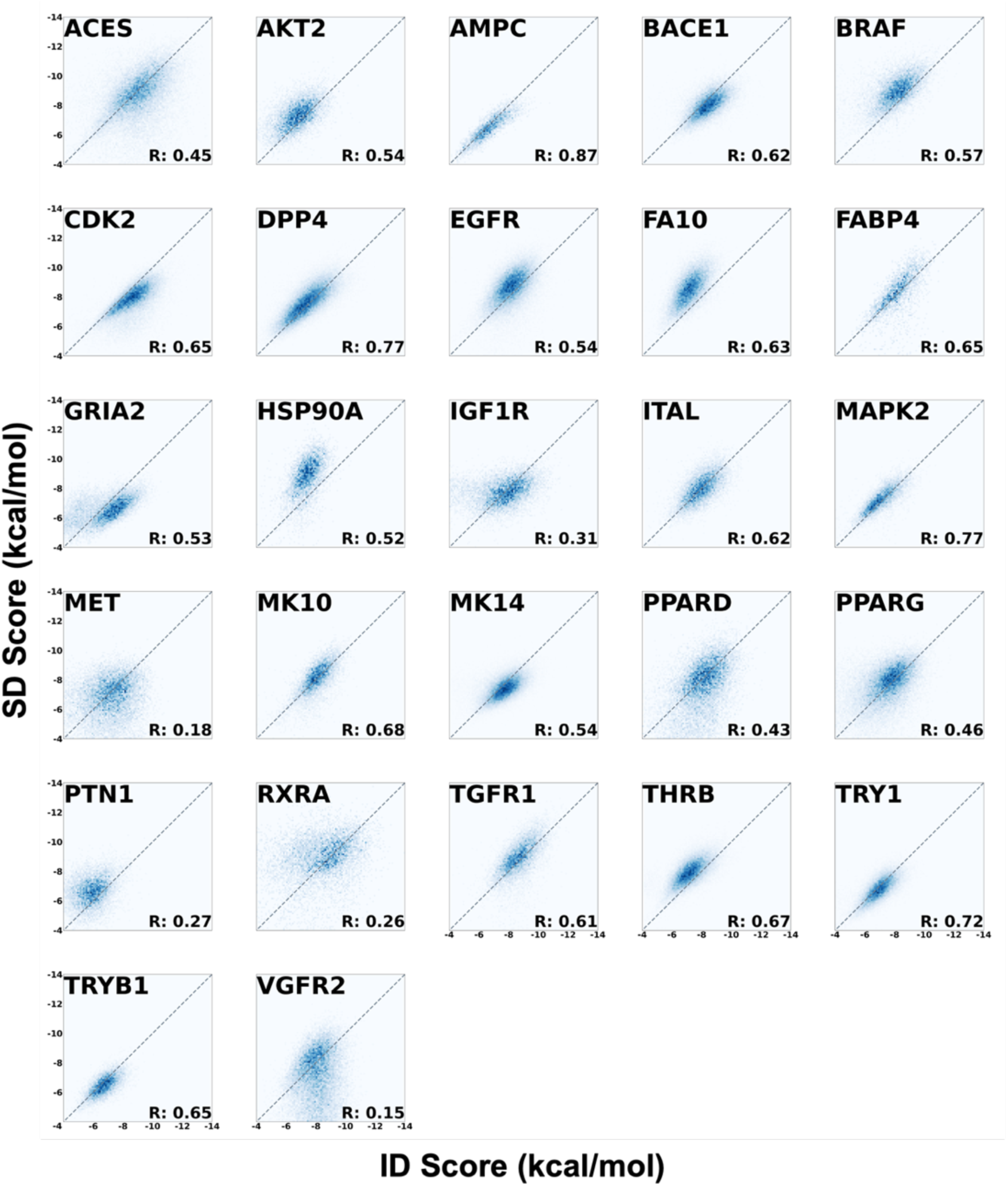
Comparison of docking scores between ID using NP and SD pipelines.

### Evaluation of consensus ranking pipelines

An important observation from our study is that individual docking is able to identify distinct active compounds compared to standard docking. The top 1% ranked compounds using ID-NP and SD also showed a low overlap rate of 15.0%. We further explore the possibility to combine results from both pipelines to further enhance the VS enrichment. Two such consensus ranking methods were evaluated: one based on docking scores and the other utilizing the Borda Count approach. In the first method, each compound was assigned the smaller of the SD and ID-NP docking scores as its final score, based on which compounds were ranked. In the second consensus ranking method, each compound was assigned a rank in both the SD and ID-NP pipelines based on its respective docking scores. The rank *m* of a compound was converted into a score of (*N*−*m*) where *N* is the total number of compounds in the VS library. This conversion ensures that the score is inversely proportional to the rank, assigning higher scores to higher-ranked compounds. The scores from both pipelines were then summed to generated a final score used for ranking.

The EFs at the 1% cutoff were calculated after re-ranking compounds using these methods (Table 1). In the docking score-based consensus ranking, nine systems achieved a higher EF than SD, four systems were identical (including two systems where both EFs were 0), and 14 systems had a lower EF compared to SD. Similarly, in the Borda Count-based consensus ranking, 10 systems outperformed SD, three systems had the same EF (including two with EF=0), and 14 systems performed worse. Notably, the median EF across the 27 systems improved for both consensus methods compared to SD, increasing slightly from 3.08 to 3.36 and 3.52, respectively. These results demonstrate that the additional information generated by individual docking could potentially be used to enhance VS workflows.

### Impact of docking on NeuralPLexer-sampled ligand poses

It’s interesting to analyze the impact of docking on NeuralPLexer-predicted ligand binding poses. Figure 5A shows the distribution of DockRMSD values with ligand kept rigid (ID-rigid-NP) or flexible (ID-NP) during the docking calculations. The average DockRMSD values were 6.41 Å and 6.47 Å for ID-NP and ID-rigid-NP, with median values of 6.40 Å and 6.60 Å, respectively. These significant deviations indicate substantial changes in ligand conformations, sometimes even resulting in reversals of binding poses within the pocket. Such results suggest that binding pose information provided by NeuralPLexer is hardly recognized or utilized by Vina, highlighting the incompatibility between DL-based predictions and physics-based docking calculations.

**Figure 5.**
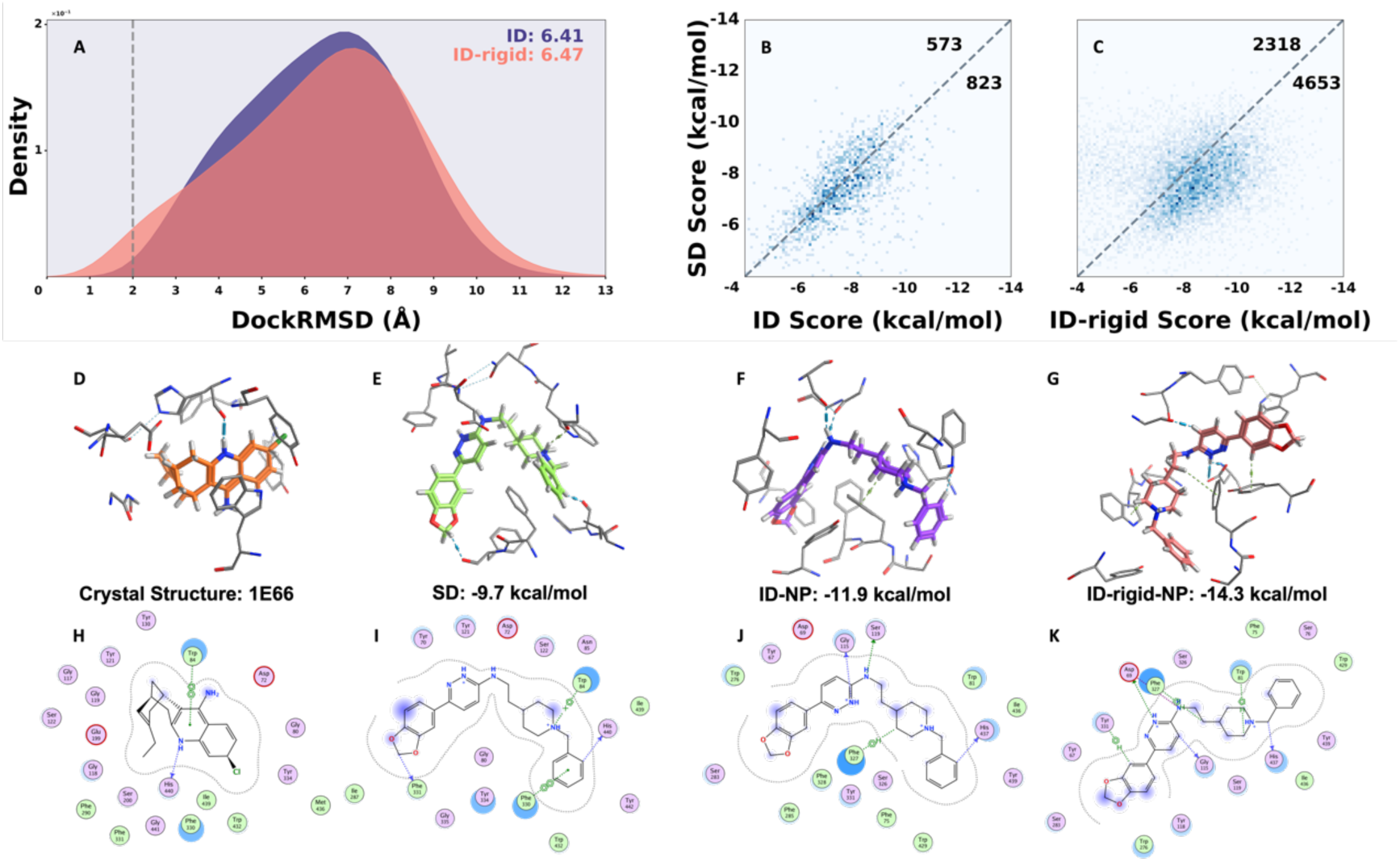
Comparison of SD, ID-NP, and ID-rigid-NP pipelines in ligand binding poses. A: Distribution of ligand DockRMSD values with the ID-NP and ID-rigid-NP pipelines. B-C: Comparison of docking scores for compounds within DockRMSD values less than 2 Å. D-G: Binding poses and interactions for crystal structure, SD, ID-NP, and ID-rigid-NP pipelines. H-K: Detailed interaction diagrams for crystal structure, SD, ID-NP, and ID-rigid-NP.

We further analyzed a minority of cases where DockRMSD values were less than 2 Å, hypothesizing that Vina might be able to effectively utilize the protein-ligand information captured by NeuralPLexer in these instances. We extracted this subset from either ID-NP or ID-rigid-NP to analyze the distribution of docking scores (Fig. 5, B and C). In the ID-NP pipeline, there were 1,396 cases with ligand RMSD less than 2 Å, of which 823 (59.0%) had docking scores lower than those from SD. ID-rigid-NP showed an increased portion, with 4,653 of 6,971 cases (66.7%) having more negative docking scores compared to SD ones.

To demonstrate how the docking scores of these cases improve through individual docking pipelines, we selected a representative case to conduct conformational analysis, as shown in Fig. 5. This case involved the docking result of the 77th active ligand (CHEMBL1202881) targeting ACES. The ligand DockRMSD values for ID-NP and ID-rigid-NP between post-sampling and post-docking stages were 1.19 Å and 0.69 Å, respectively, indicating that both pipelines successfully preserved the deep model’s understanding of the ligand binding pose. Consequently, the conformations in SD, ID-NP, and ID-Rigid-NP reflected an increasing trend in retaining information from the DL model. A comparison with the crystal binding mode (Fig. 5, D and H) further highlighted the advantages of DL model sampling in improving docking scores.

In SD, which did not contain any NeuralPLexer information (Fig. 5, E and I), the ligand’s right-terminal phenyl group appeared in a position similar to that in the crystal structure and successfully reproduced the π-π stacking interaction with W84 and hydrogen bonding with H440. However, the larger SD ligand was forced to fit the protein pocket in an over-bending conformation, consistent with the observations from the ACES 22,573rd decoy case (Fig. 3C). In the ID-NP binding mode, the protein pocket underwent significant conformational changes (PPS score: 0.52, pocket RMSD: 1.19 Å) induced by the ligand, allowing the ligand to adopt a conformation closer to its natural state. This adjustment also revealed that this larger ligand utilized its pyridazine group and aromatic amine to form new hydrogen bonds with G118 and S122, enhancing ligand-protein interactions. However, ID-NP failed to recognize W84 as a key interacting residue, which is crucial for real binding and a desirable feature to be reproduced in molecular docking.

These limitations observed in SD and ID-NP were effectively optimized in this case by ID-rigid-NP (Fig. 5, G and K). By fully retaining the DL-sampled information on both the protein pocket and ligand conformation, ID-rigid-NP generated an energetically very favorable ligand conformation. It also successfully reproduced the key contributions of W84 and H440 to the binding mode. Moreover, ID-rigid-NP identified additional interactions, including backbone and side-chain hydrogen bonds with the 1,3-benzodioxole and pyridazine groups of the ligand compared. Overall, these observations align with the trends noted in the conformational analysis of the ACES decoy ligand: DL models tend to sample protein pockets and ligand conformations that maximize ligand-protein interactions, which might also contribute to enhanced docking scores in some cases.

Subsequently, we calculated the EFs of these two subsets. In the ID-NP subset, among 1396 ligands, 60 were active, yielding an equivalent enrichment factor of 1.67, which remains below the average level. Similarly, in the ID-rigid-NP subset, no improvement in VS performance was observed, with an equivalent EF of 1.66 or 297 active hits out of 6,971 cases. This outcome aligns with our previous observation: even when the conformational sampling performed by the DL model can optimize docking scores for protein-ligand pairs (in the ID-rigid-NP subset, the proportion is 2/3), it does not make active ligands bind more favorably relative stronger to decoy compounds. Instead, it may represent a non-discriminatory fitting approach that universally forms more ligand-protein interactions.

### The VS performance for individual docking by AlphaFold3

We further applied the individual docking protocol using AlphaFold3 as the structure prediction method. As with NeuralPlexer, we considered three docking protocols: ID-scoring (scoring predicted complexes), ID-rigid (rigid docking into predicted structures), and ID (flexible ligand docking), with Vina used as the docking engine. The ID-scoring-AF3 pipeline yielded 54,505 complexes (12.23%) with docking scores >0 kcal/mol, a significant reduction compared to ID-scoring-NP (96.42%). This lower incidence of positive scores suggests that AF3’s structure predictions more effectively minimize steric clashes, rendering direct scoring a viable option for virtual screening.

Individual docking with AF3 demonstrated consistent advantages over standard docking. As summarized in Table 2, S2, and S3, all three AF3-based ID pipelines showed improved enrichment factors at the 1% cutoff, with mean values of 12.45, 9.51, and 7.10 (medians: 10.22, 7.23, and 6.18), respectively. ID-scoring-AF3 demonstrated particularly strong performance, achieving a 2.17-fold improvement in EF over SD and outperforming SD in 20 out of 27 systems. Notably, it successfully identified an active compound in AMPC, where SD failed (Table 2). However, reduced performance was observed in five systems (e.g., AKT2 where SD identified active ligands while ID-scoring-AF3 failed), and two systems showed equivalent performance (DPP4 and HSP90A). Further analysis demonstrated that docking score-based consensus rankings produced highly similar EF values across AF3 pipelines (Table 2, S2, and S3). The superior discriminative capability of ID-scoring-AF3 was also evident in its average AUC of 0.75, with outstanding performance (AUC>0.9) in BACE1, BRAF, MK14, and RXRA (Fig. 6). Comparative performance across all systems is provided in Fig. S3, illustrating both the strengths and limitations of the AF3-based approaches.

**Table 2.** Virtual screening performance of ID-scoring with AF3 evaluated at 1% cutoff.

| System | Enrichment Factor |  | # Active Hits<br>(SD/ID/Shared) | Overlap Rate | Consensus Ranking |  |
| --- | --- | --- | --- | --- | --- | --- |
|  | SD | ID-scoring |  |  | Docking Score-Based | Borda Count-Based |
| ACES | 3.31 | 18.97 | 15/86/6 | 4.5% | 18.97 | 12.36 |
| AKT2 | 1.69 | 0.00 | 2/0/0 | 1.4% | 0.0 | 4.22 |
| AMPC | 0.00 | 1.59 | 0/1/0 | 3.3% | 1.59 | 1.59 |
| BACE1 | 4.51 | 36.52 | 22/178/11 | 5.9% | 36.52 | 21.75 |
| BRAF | 15.46 | 24.98 | 39/63/18 | 18.3% | 24.98 | 27.36 |
| CDK2 | 8.62 | 3.50 | 69/28/13 | 8.2% | 3.50 | 12.62 |
| DPP4 | 0.74 | 0.74 | 8/8/2 | 16.7% | 0.65 | 1.20 |
| EGFR | 3.36 | 4.44 | 28/37/1 | 1.7% | 3.84 | 6.61 |
| FA10 | 8.17 | 23.01 | 65/183/50 | 24.9% | 23.01 | 16.97 |
| FABP4 | 18.73 | 10.22 | 11/6/5 | 20.0% | 10.22 | 28.95 |
| GRIA2 | 3.02 | 2.68 | 9/8/0 | 4.8% | 2.68 | 5.37 |
| HSP90A | 0.00 | 0.00 | 0/0/0 | 0.0% | 0.00 | 0.00 |
| IGF1R | 3.08 | 7.91 | 7/18/1 | 2.1% | 7.91 | 11.86 |
| ITAL | 0.43 | 15.32 | 1/36/1 | 1.1% | 15.32 | 6.38 |
| MAPK2 | 6.74 | 11.56 | 14/24/2 | 6.2% | 11.56 | 12.04 |
| MET | 1.64 | 10.63 | 4/26/2 | 1.7% | 10.63 | 10.63 |
| MK10 | 1.59 | 14.31 | 3/27/2 | 10.0% | 14.31 | 9.01 |
| MK14 | 1.64 | 24.01 | 15/220/3 | 1.3% | 24.01 | 10.26 |
| PPARD | 2.42 | 7.94 | 7/23/0 | 2.2% | 7.94 | 7.94 |
| PPARG | 5.12 | 12.44 | 37/90/19 | 9.4% | 12.44 | 12.86 |
| PTN1 | 7.07 | 7.96 | 16/18/0 | 5.2% | 7.96 | 11.49 |
| RXRA | 28.90 | 34.43 | 47/56/24 | 30.4% | 34.43 | 40.58 |
| TGFR1 | 11.69 | 7.44 | 33/21/1 | 3.3% | 7.44 | 10.27 |
| THRB | 3.60 | 7.54 | 31/65/5 | 8.2% | 7.54 | 8.12 |
| TRY1 | 2.24 | 3.42 | 17/26/0 | 3.0% | 3.42 | 3.95 |
| TRYB1 | 1.17 | 16.92 | 2/29/1 | 1.3% | 16.92 | 5.84 |
| VGFR2 | 2.73 | 27.80 | 17/173/6 | 2.7% | 27.80 | 6.27 |
| **AVERAGE** | **5.74** | **12.45** | -- | 7.3% | **12.43** | **11.35** |
| **MEDIAN** | **3.08** | **10.22** | -- | 4.5% | **10.22** | **10.26** |

**Figure 6.**
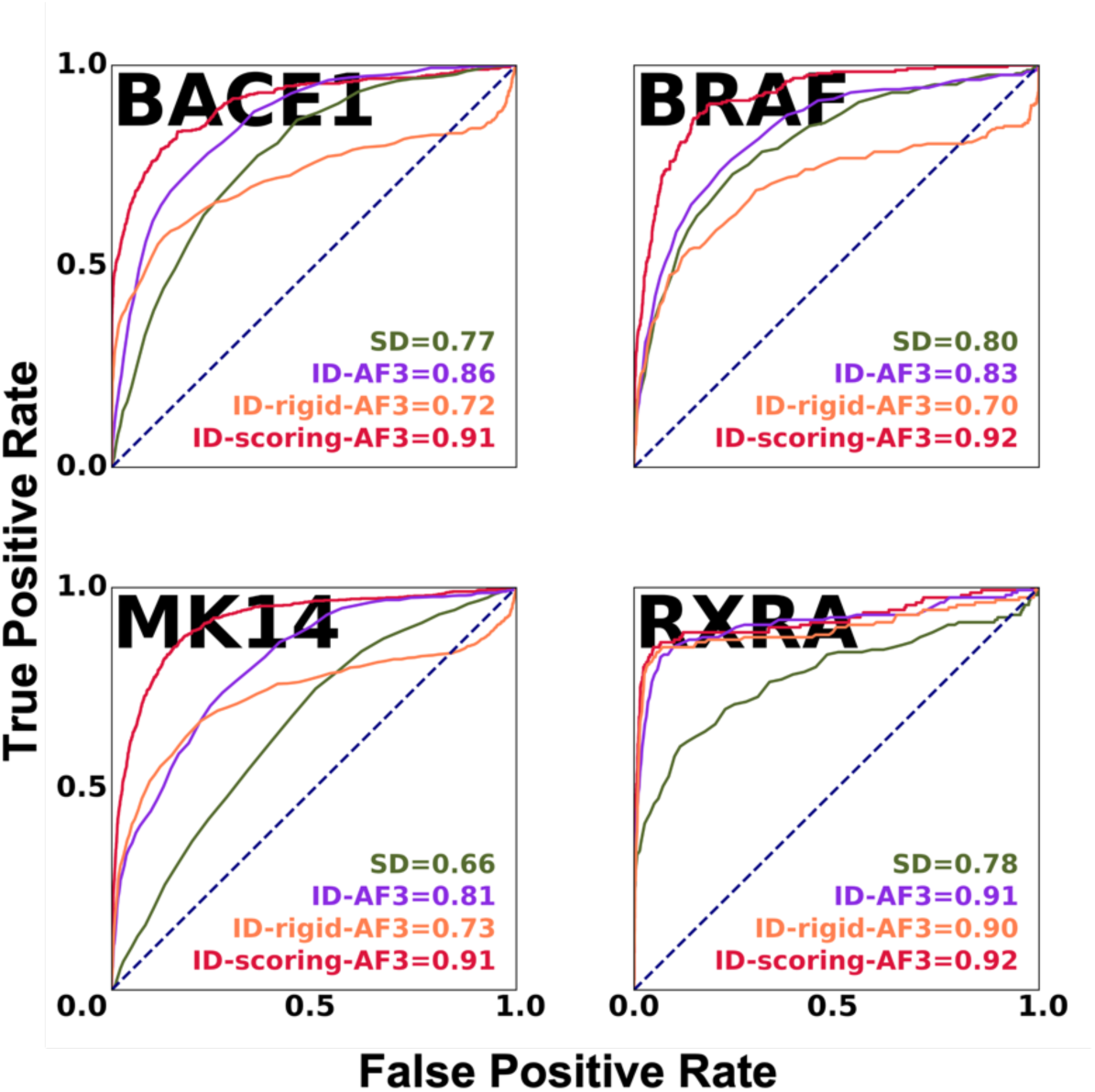
ROC curves and AUC values for BACE1, BRAF, MK14 and RXRA systems sampled by AF3, in which VS performance has been significantly improved with individual docking.

Similar to the NeuralPLexer-based results, the AF3-based docking pipelines exhibited remarkably low overlap with SD in their ligand selections. Quantitative analysis revealed average top 1% overlap rates of just 7.3% (ID-scoring-AF3), 8.5% (ID-rigid-AF3), and 14.0% (ID-AF3) with SD, with corresponding median values of 4.5%, 6.0%, and 12.5% (Table 2, S2, and S3). This divergence was even more pronounced for active compounds, suggesting that AF3-based individual docking and SD often identify distinct sets of bioactive molecules. Notably, in systems like EGFR, PTN1, GRIA2, IGF1R, and TRY1, where both methods showed comparable enrichment performance, the number of shared active hits remained below one in all cases. Interestingly, consensus ranking via Borda Count improved EF1% by approximately two-fold, highlighting the complementary nature of ID and SD approaches.

### AlphaFold3 yields discriminatory power between actives and decoys

The docking score distributions for active and decoy ligands across all pipelines (ID-AF3, ID-rigid-AF3, ID-scoring-AF3, and SD) were systematically compared (Fig. S4). Both SD and ID-AF3 exhibited narrow, peaked distributions with scores concentrated in the moderate range (-9 to -6 kcal/mol). In contrast, ID-rigid-AF3 and ID-scoring-AF3 displayed broader, flatter distributions, with pronounced tails in both high- and low-scoring regions. The mean docking scores were -7.60 (SD), -7.89 (ID-AF3), -8.07 (ID-rigid-AF3), and 4.17 kcal/mol (ID-scoring-AF3). Notably, despite sharing similar DL-derived structural inputs, ID-rigid-AF3 and ID-scoring-AF3 yielded the most divergent mean scores, suggesting distinct scoring behaviors.

Further stratification by ligand type (Fig. 7, A and B) showed that active ligands in ID-rigid-AF3 and ID-scoring-AF3 had similar distributions, both skewed toward more favorable (lower) scores compared to SD or ID-AF3. However, decoy ligands in ID-scoring-AF3 exhibited a unique shift toward less favorable (higher) scores, resulting in a clear separation between actives (mean: -8.63 kcal/mol) and decoys (mean: -4.05 kcal/mol). This enhanced discriminative ability aligns well with the superior VS performance of ID-scoring-AF3. Such a fundamental improvement in distinguishing actives within AF3-predicted complexes, as reflected by the distinct docking score distributions, can be attributed to the increased accuracy and precision in modeling protein-ligand interactions.

**Figure 7.**
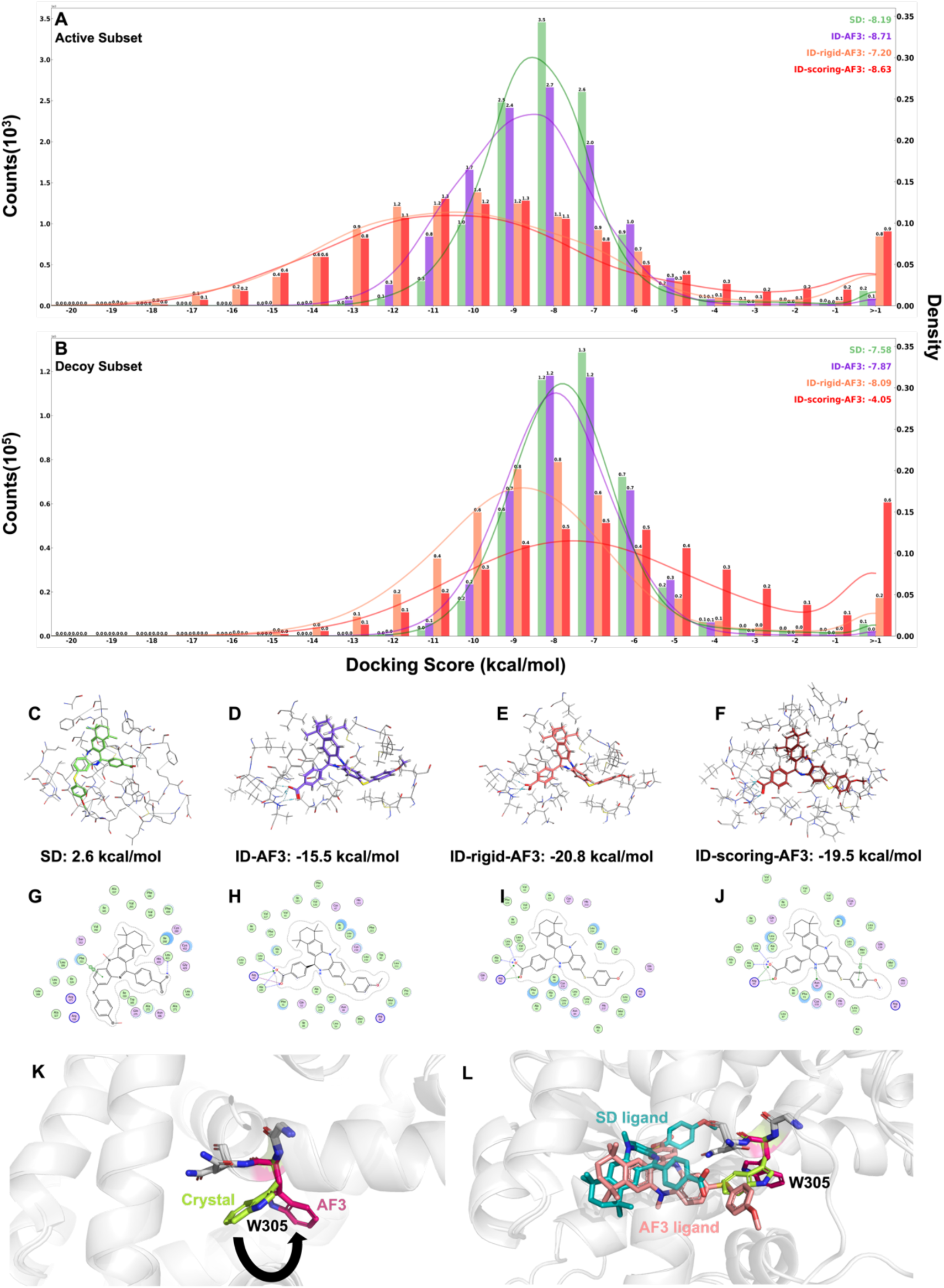
Comparison of docking score distribution of SD, ID-AF3, ID-rigid-AF3, and ID-scoring-AF3 pipelines. A: Docking score distribution of active ligands. B: Docking score distribution of decoy ligands. C-F: Binding poses and interactions of each pipeline for ligand CHEMBL396658 targeting RXRA. G-J: Detailed interaction diagrams for each binding pose. K: Conformation change of W305 from the crystal structure (green) to the AF3-predicted structure (purple). L: Binding pose comparison between SD (blue) and ID-scoring-AF3 (pink).

As an example, the RXRA-CHEMBL396658 complex achieved very favorable docking scores in both ID-scoring-AF3 (-19.5 kcal/mol) and ID-rigid-AF3 (-20.8 kcal/mol). AF3’s predicted binding pose (Fig. 7, F and J) revealed an extensive interaction network, including hydrogen bonds with N84, R94, L104, and A105, plus aromatic interactions with M239. While ID-rigid-AF3 and ID-AF3 reproduced key interactions (Fig. 7, H and I), SD failed completely, yielding a positive score (2.6 kcal/mol) with only two weak interactions (Fig. 7, C and G). This improvement stems from AF3’s prediction of an outward rotation of W305 (Fig. 7K), shifting its dihedral angle from -58° (gauche-) to 69° (gauche+), which substantially enlarges the binding pocket to accommodate the bulky ligand (Fig. 7L). Notably, re-docking into this enlarged pocket did not fully recover the key interactions, the ID-AF3 docking score was -15.5 kcal/mol, further underscoring the inherent incompatibility between DL-generated pocket conformations and conventional docking algorithms.

These findings collectively demonstrate the transformative potential of individual docking. By generating dynamic, ligand-adapted binding pocket conformations through DL-based approaches, it is possible to circumvent the spatial limitations imposed by static crystal structures. This adaptive capability is particularly valuable for challenging ligands like CHEMBL396658, effectively overcoming one of the longstanding obstacles in structure-based drug discovery.

## Discussion and Conclusions

The flexibility of protein receptor conformations is crucial for accurately predicting potential binding ligands. For example, Mobley and co-workers showed in a model system (13 ligands targeting the L99A mutant of T4 lysozyme) that performing energy minimization of ligand-bound proteins significantly improved the correlation between calculated absolute binding free energies and experimental values, increasing the correlation coefficient from -0.05 to 0.82 [52]. DL-based structure prediction methods, such as NeuralPLexer and AlphaFold3, provide a new and computationally efficient way to generate ligand-specific protein structures. The large-scale generation of ligand-protein complex structures creates an opportunity to mitigate the impact of assuming a fixed receptor conformation during docking. This study explored the concept of individual docking, in which complex structures were predicted for each individual ligand using DL models and the predicted protein structures were used for docking calculations. While showing the feasibility of ID, our findings reveal several critical limitations that hinder its VS performance.

We performed systematic benchmarking of individual docking workflows versus standard docking protocols across 27 DUD-E targets. Our results show that VS performance is highly sensitive to the choice of DL models for conformational sampling. While NP-based ID workflows showed consistently inferior performance with both Vina and MOE docking methods (Table S4, Figure S5), AF3-based ID pipelines achieved substantially improved screening outcomes. Analysis of docking score distributions suggests that AF3 possesses an inherent ability to distinguish active compounds from decoys. However, this discriminative power was significantly compromised when docking was applied to the AF3-predicted structures, as evidenced by the reduced EFs observed in both ID-AF3 and ID-rigid-AF3 pipelines compared to ID-scoring-AF3

An in-depth analysis of pocket conformational changes using the PPS score demonstrated that NeuralPLexer is capable to generate pockets geometrically tailored to individual ligands, a unique capability not present in traditional VS approaches. However, this customization comes at the cost of reduced compatibility with the energetics (empirical docking scores) used for scoring and sampling in docking workflows. Variations in docking score distributions among SD, ID-NP, and ID-rigid-NP pipelines suggest that the DL model struggles to accurately model the subtle physical interactions essential for protein-ligand binding. Additionally, its predictions failed to reliably distinguish between actives and decoys, as evidenced by specific examples.

This limitation might be partly due to NeuralPLexer prioritizing spatial arrangements based on distance constraints rather than the physical and chemical properties critical to binding interactions. Further analysis of the DockRMSDs between NeuralPLexer-generated poses and docking-refined poses revealed significant rearrangements of the ligand from the predicted structure, addressing atomic clashes and enhancing interactions between protein and ligand atoms according to docking energy evaluations.

In addition, structural analysis revealed substantial conformational differences between AF3-generated poses and those produced by subsequent docking. DockRMSD measurements showed average deviations of 4.89 Å (median: 4.96 Å) for ID-AF3 and 3.09 Å (median: 1.22 Å) for ID-rigid-AF3 relative to the original AF3 predictions (Fig. S6). Notably, these deviations showed similar distributions for both active and decoy compounds (Fig. S7). Interestingly, the AF3-generated pockets exhibited higher similarity to crystal structures (mean PPS score: 0.68; pocket RMSD: 0.55 Å). These subtle structural perturbations, induced by AF3’s sampling of binding pocket conformations, appear to shift the docking score in a less discriminatory way between active and decoy ligands during re-docking, consequently impairing enrichment performance relative to direct scoring on AF3-generated poses.

We note that out study employed two popular docking programs (Vina and MOE), but the performance of individual docking may vary with more sophisticated computational approaches, such as DL-based scoring functions [53,54], induced-fit docking [55], or free energy perturbation (FEP) calculations [7]. In particular, simulation-based FEP methods, such as the commercially available FEP+ software [56], could potentially improve accuracy by better modeling protein flexibility and binding thermodynamics. This will be further explored in future work.

To improve the performance of AI-based ID, two key strategies hold promise. First, incorporating physical interaction constraints during AI model training could improve the structural realism and interpretability of predicted protein-ligand complexes, making them more compatible with physics-based binding free energy estimations [57]. Second, training or fine-tuning models to distinguish actives from decoys might increase the accuracy and reliability of structure predictions by injecting deeper insights into ligand-protein interactions. We also note that the landscape of ligand-protein complex prediction models is advancing rapidly. For example, NeuralPLexer, the conformational sampling model used in this study, has already progressed to its third-generation version [41]. Alternative implementations of AlphaFold3 have also been emerged, including Protenix [58], Chai-1 [42] and Boltz-1 [43]. Although the computational cost associated with generating individual pockets remains relatively high, the observed improvement in VS performance suggests that individual docking could serve as a valuable complement to standard docking approaches. For example, ID might be particularly useful for prioritizing hits from DNA-encoded library (DEL) screening. The insights obtained in this study lay a foundation for more effective integration of DL-based and physics-based VS methods, advancing computational drug discovery.

## Methods

### Dataset

This study employs a subset of the DUD-E dataset, specifically DUD-E 27, as the benchmark for individual docking experiments. The dataset includes 27 protein targets and a total of 445,730 compounds, comprising 11,447 active compounds and 434,283 decoy compounds. All ligand data were downloaded from the official DUD-E website (https://dude.docking.org/) and formatted using Molecular Operating Environment (MOE2022) software [59]. Additionally, 3D structures for all proteins were retrieved in PDB format from the Protein Data Bank (PDB) website (https://www.rcsb.org/), in accordance with the descriptions provided by DUD-E.

### Protocol of standard docking

For standard docking, both Vina and MOE [59] were used with a semi-flexible docking protocal. In the Vina workflow, receptor and ligand files were converted from PDB to PDBQT format using MGLTools [60], during which the receptor conformations were fixed while the ligand ones were set flexible. Docking site was defined as a cube with a side length of 15 Å, with the center of mass of the ligand in each crystal structure serving as the central point. In the MOE docking setup, energy minimizations were performed to prepare the protein 3D structures for docking. The docking site was automatically defined by MOE through selecting the ‘ligand’ option, in which MOE would detect the space within 5Å of ligand atoms to guide the placement. ‘Rigid receptor’ was selected to apply the semi-flexible docking approach. Parameters without precise explanations, such as scoring functions, placement algorithms, were set as default.

By default Vina outputs top five docking poses (30 poses in MOE approach) after conformation search. The pose with the best docking score (numerically lowest) was selected for further analysis.

### Protocol of individual docking

During the NP sampling process, the ligand in SDF format and the receptor in PDB format were used as inputs for NeuralPLexer. The ‘batched_structure_sampling’ function of NeuralPLexer [40] predicted 16 complex structures, with the one having the highest pLDDT score selected as the final result. Templates were not used during the sampling process to prevent generating receptor conformations with over-influenced by the templates, thus enabling ligand-induced conformational fitting. In the process of AF3, protein sequence and ligand smiles were written into json file as input for AF3. The complex structure with highest pLDDT was selected for subsequent docking. The inference process is conducted in parallel, with each parallel execution taking approximately six hours for NP and nine hours for AF3 to predict about 1,000 complex structures on an NVIDIA L40 GPU core. All unspecified parameters were set to their default values.

In the docking process, the ID-scoring pipelines utilize the ‘--score_only’ option in the Vina program, in which only protein and ligand PDBQT files were input. In the ID pipelines, semi-flexible docking was applied by setting the ‘inactivate_all_torsions’ option to ‘False’ during the processing step of ligand files. In contrast, in the ID-rigid pipelines, rigid docking was applied by setting this option activated, to preserve the ligand in the same pose as predicted by NeuralPLexer. Other parameter setup kept the same as SD.

### Assessment of pocket similarity and ligand RMSD

We use PPS-align [50] to assess the conformational differences between each sampled pocket predicted by NP/AF3 and its corresponding crystal structure. Residues within 5 Å of the ligand atoms are defined as the pocket. DockRMSD [51] is then used to evaluate the ligand RMSD between the NP/AF3 predictions and the post-docking results from both the ID and ID-rigid pipelines. This analysis aims to compare the ligand pose differences between DL predictions and the results after docking.

## Acknowledgments

The work is supported by the “Pioneer” and “Leading Goose” R&D Program of Zhejiang (2023C03109, 2024SSYS0036), the National Natural Science Foundation of China (32171247), the Zhejiang Provincial Natural Science Foundation of China (LQ23F020011), and the Westlake Education Foundation. We thank the Westlake University Supercomputer Center for computational resources and related assistance.

