## Supplemental Materials for "Scrutinization on Docking Against Individually Generated Target Pockets for Each Ligand"

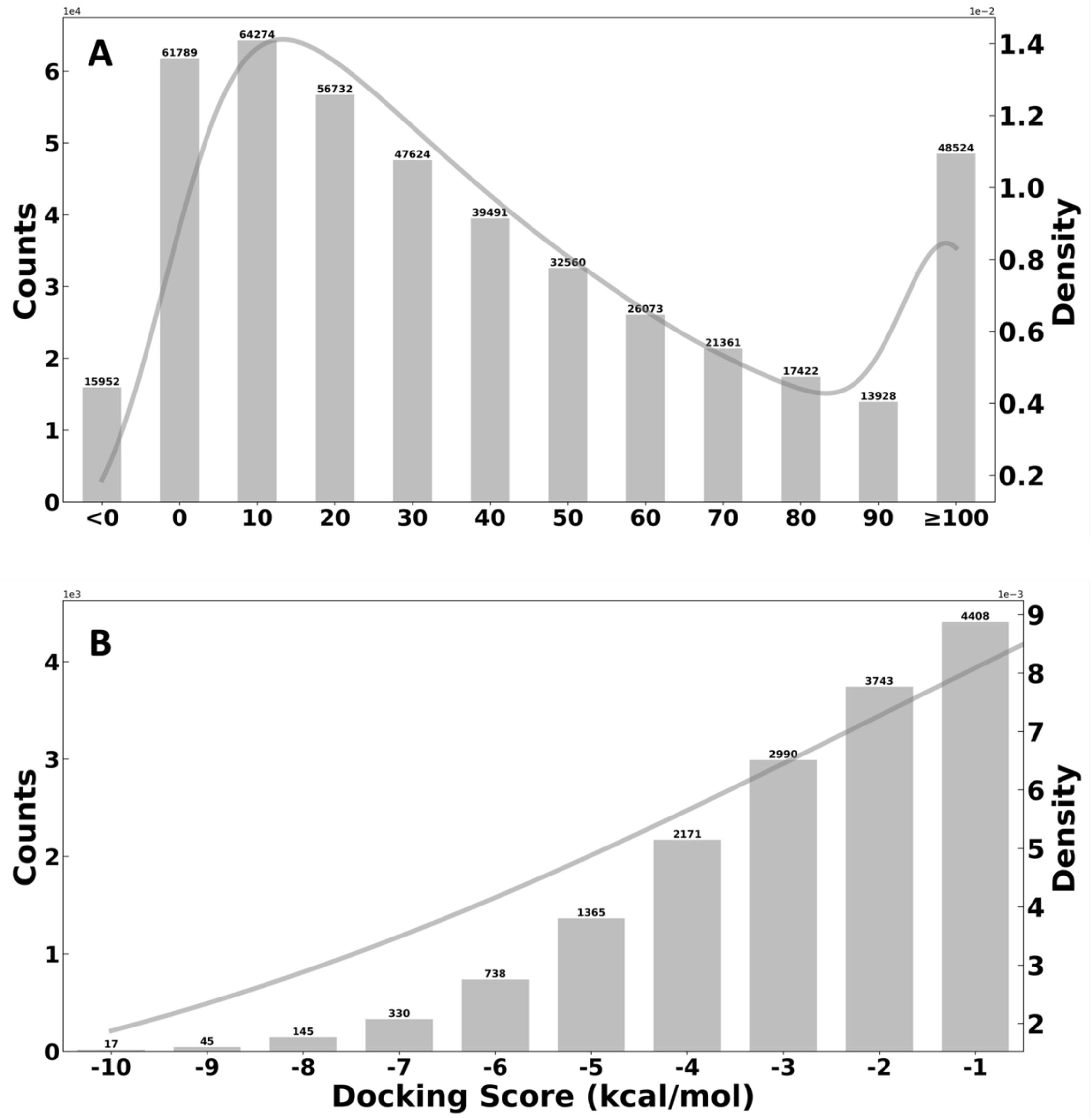


**Figure S1**. Distribution of docking score directly calculated using NeuralPLexer Structures. A: Overall docking score distribution. B: Docking score distribution for scores below 0 kcal/mol.

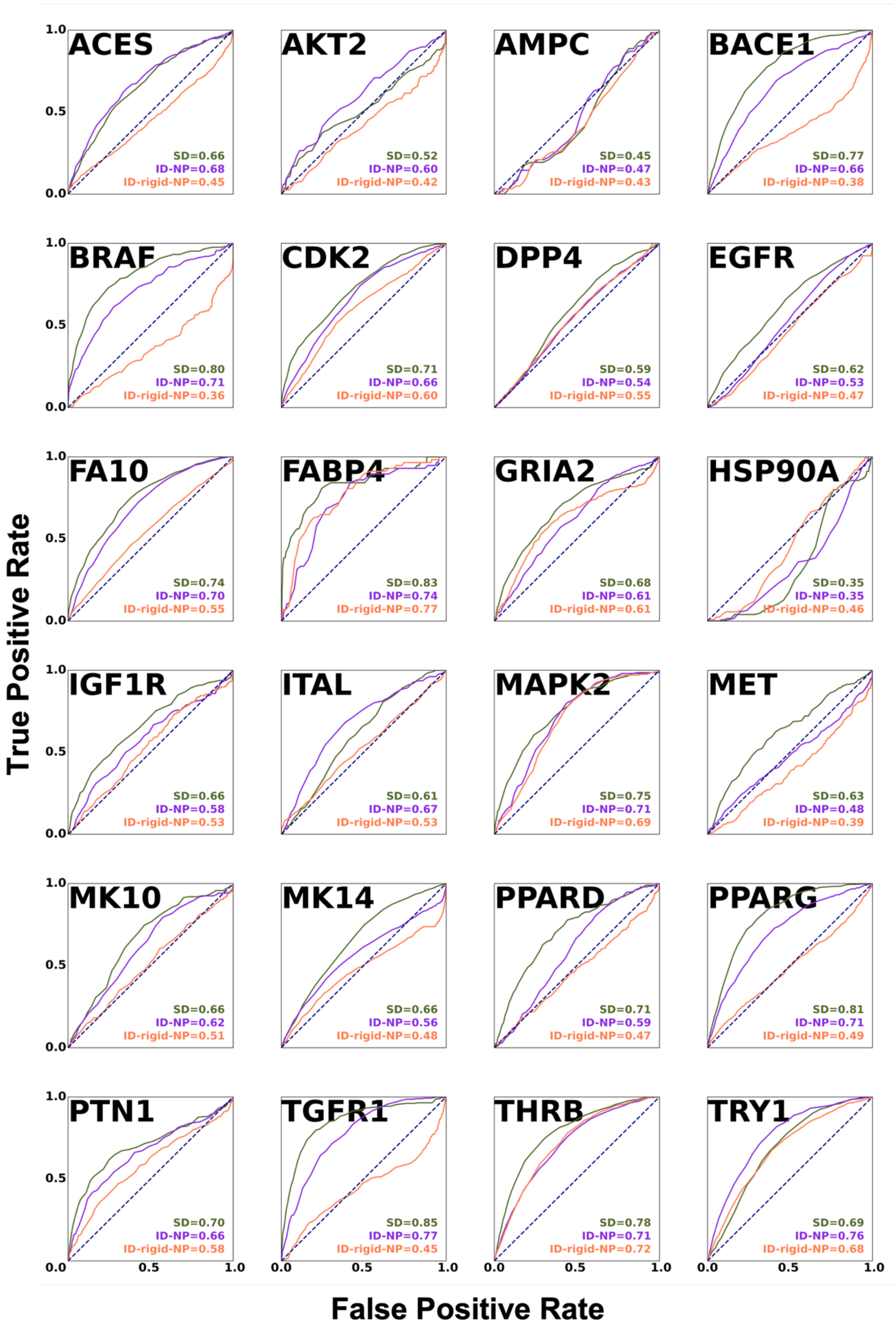


**Figure S2**. ROC curves and AUC values of 24 systems for SD, ID-NP, and ID-rigid-NP pipelines.

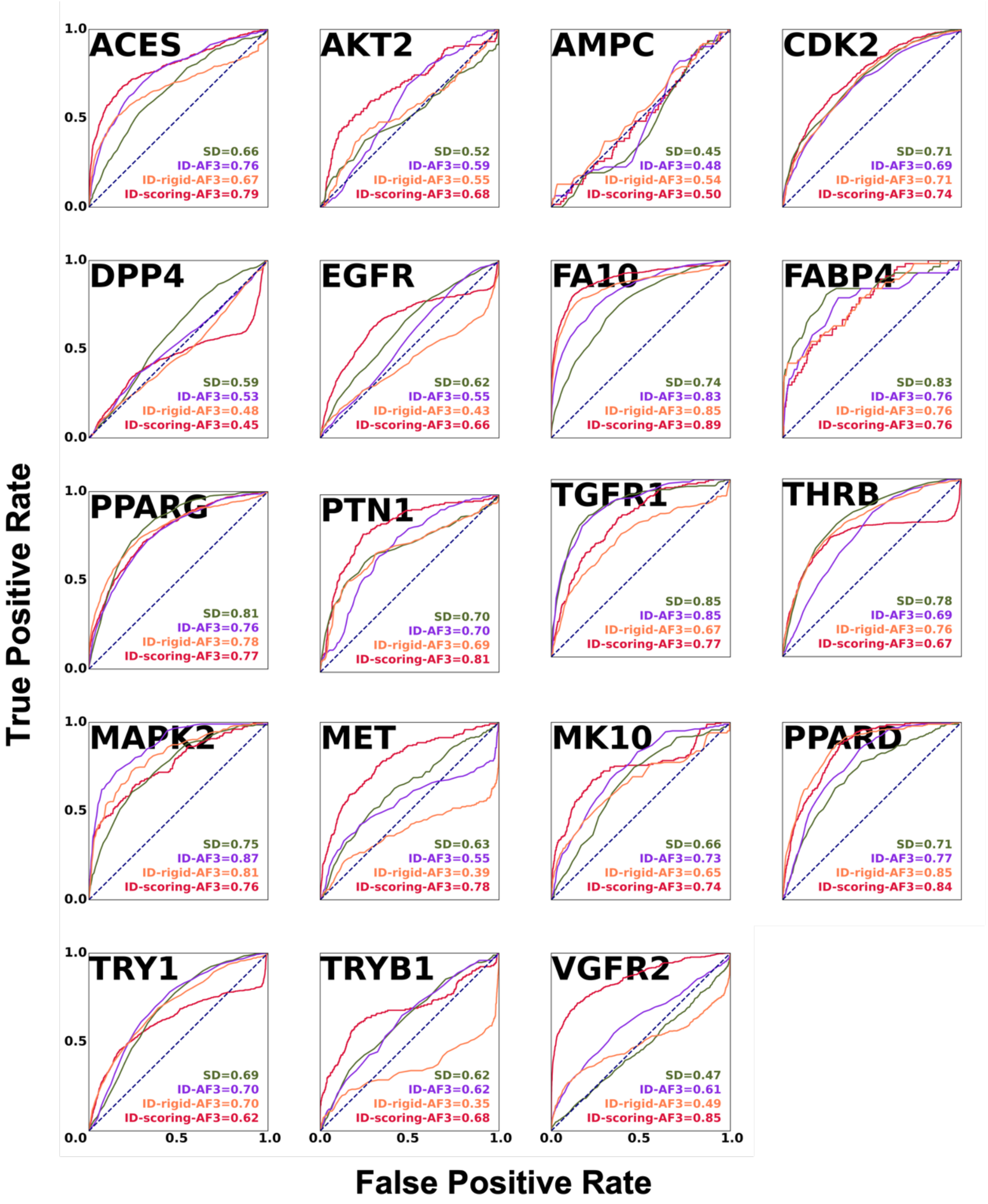


**Figure S3**. ROC curves and AUC values of 23 systems for SD, ID-AF3, ID-rigid-AF3 and ID-scoring-AF3 pipelines, respectively.

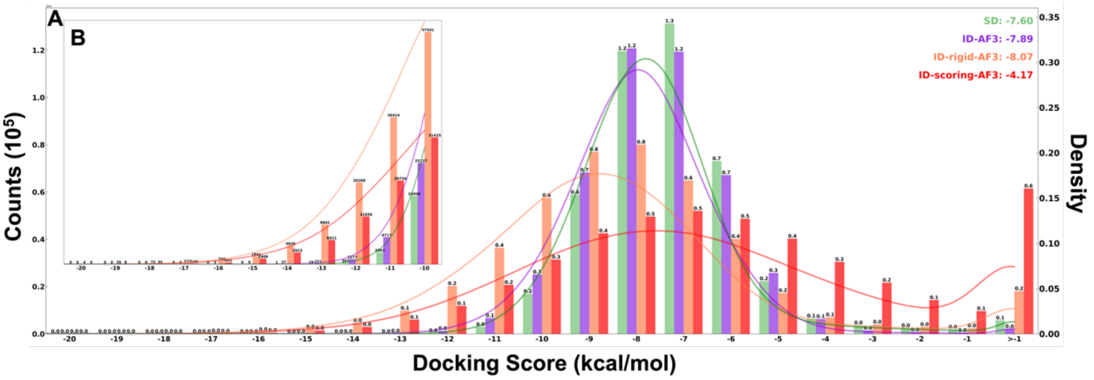


**Figure S4**. Overall docking score distributions of SD, ID-AF3, ID-rigid-AF3, and ID-scoring-AF3, respectively.

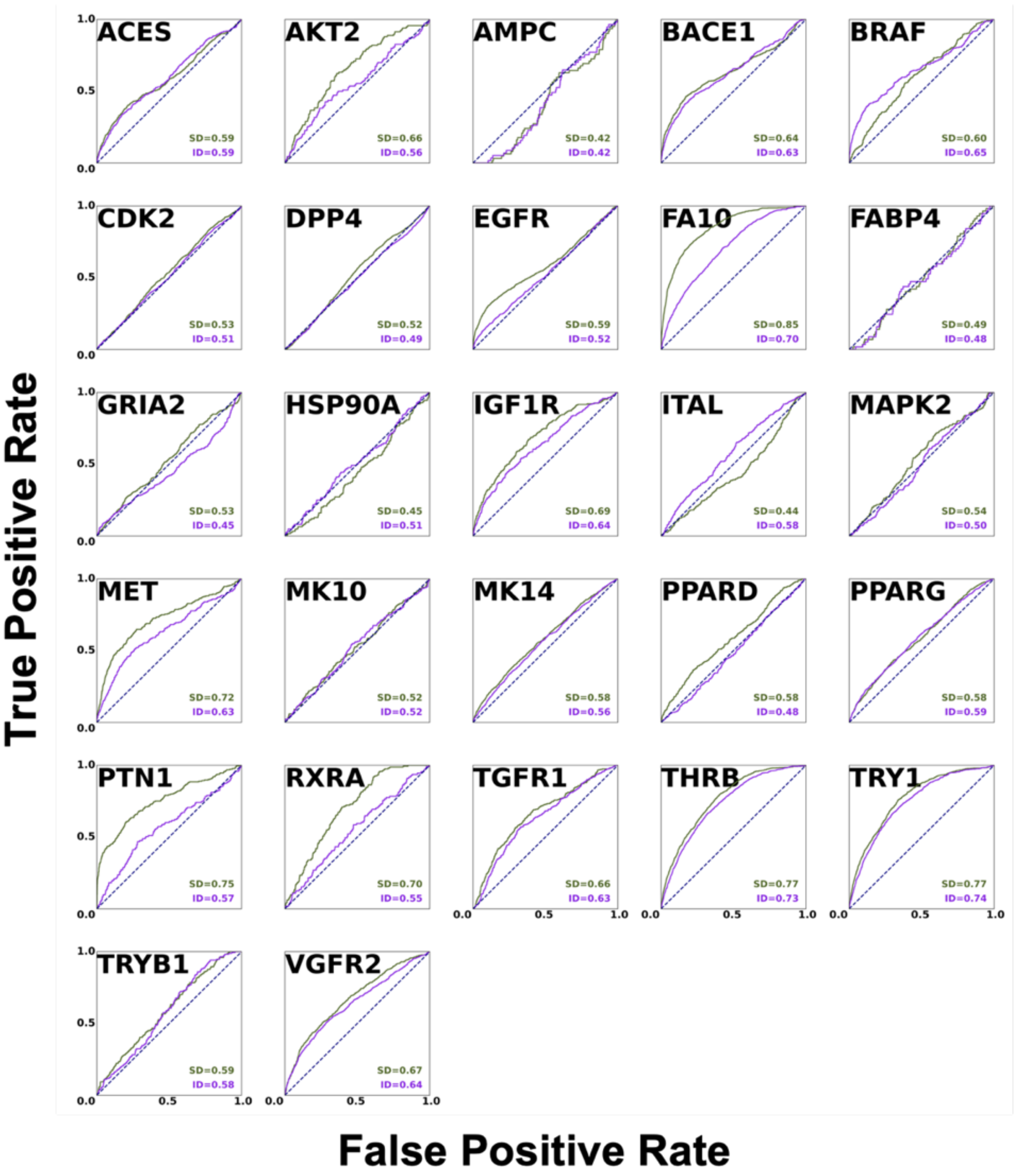


**Figure S5**. ROC curves and AUC values for standard and ID-NP pipelines utilizing MOE.

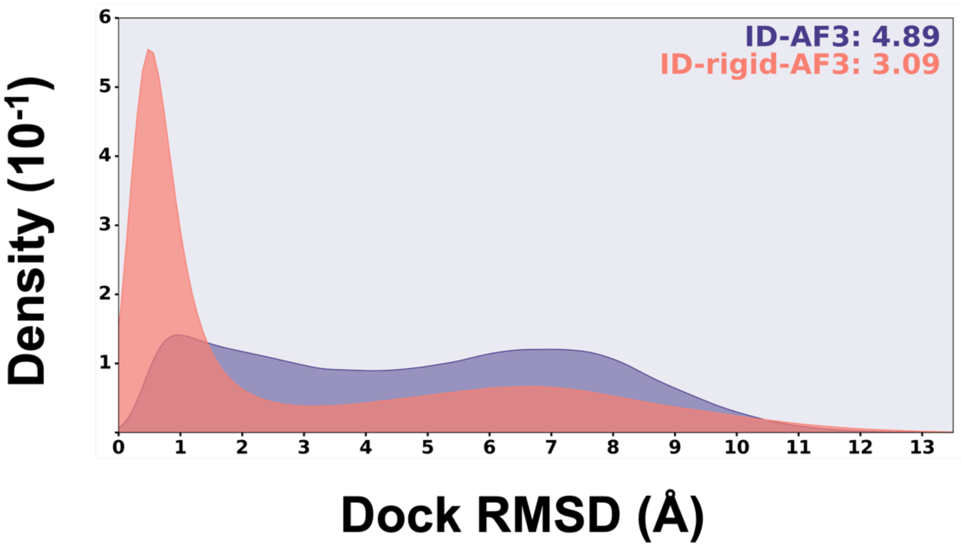


**Figure S6**. Overall dock RMSD values distributions in ID-AF3 and ID-rigid-AF3 pipelines.

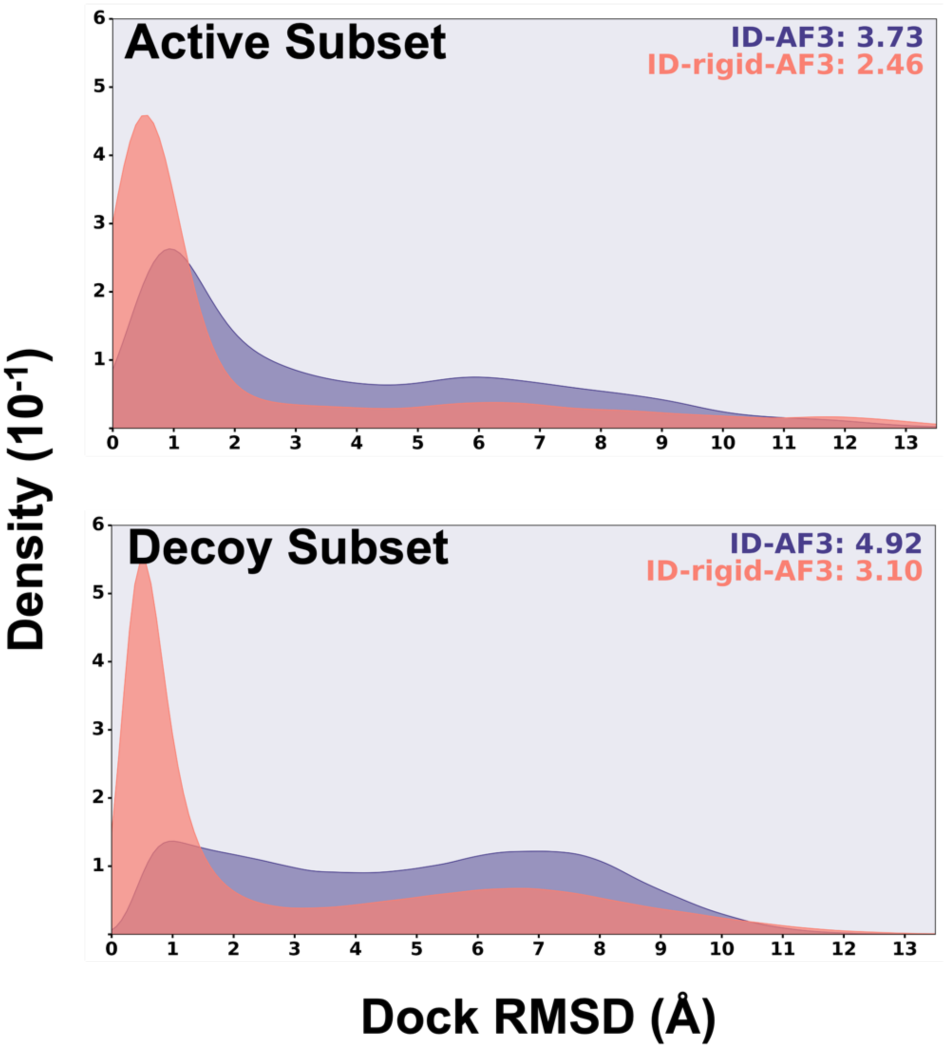


**Figure S7**. Dock RMSD values distributions of active and decoy subsets in ID-AF3 and ID-rigid-AF3 pipelines.

**Table S1. Virtual screening performance of ID-NP evaluated at 5%/10% cutoff.**

| System | SD | ID | Dock Score-based Consensus Ranking |
|---|---|---|---|
| ACES | 2.56/2.34 | 3.31/2.60 | 2.91/2.56 |
| AKT2 | 2.56/1.71 | 2.05/2.30 | 3.24/2.13 |
| AMPC | 0.00/0.48 | 0.64/0.48 | 0.64/0.32 |
| BACE1 | 3.58/3.30 | 2.02/2.21 | 2.39/2.43 |
| BRAF | 6.61/4.82 | 4.62/3.07 | 6.37/4.58 |
| CDK2 | 4.76/3.37 | 2.53/1.94 | 2.81/2.23 |
| DPP4 | 1.11/1.15 | 0.98/0.95 | 1.02/1.11 |
| EGFR | 2.21/2.13 | 0.43/0.68 | 1.92/1.86 |
| FA10 | 4.24/3.24 | 2.65/2.59 | 4.27/3.28 |
| FABP4 | 9.10/5.42 | 2.45/3.15 | 8.75/5.42 |
| GRIA2 | 3.50/2.90 | 1.68/1.41 | 1.68/1.51 |
| HSP90A | 0.00/0.08 | 0.00/0.00 | 0.00/0.08 |
| IGF1R | 3.71/2.65 | 1.77/1.68 | 1.59/2.21 |
| ITAL | 1.03/1.24 | 2.06/2.06 | 1.71/1.93 |
| MAPK2 | 3.68/3.44 | 2.23/1.70 | 3.20/2.86 |
| MET | 2.13/2.17 | 1.39/1.43 | 1.23/1.23 |
| MK10 | 2.25/1.77 | 1.61/1.66 | 2.04/1.72 |
| MK14 | 2.10/1.99 | 2.16/1.99 | 2.29/2.27 |
| PPARD | 2.77/2.81 | 0.62/0.90 | 1.66/1.70 |
| PPARG | 3.87/3.61 | 3.35/3.00 | 3.35/3.17 |
| PTN1 | 5.60/3.95 | 3.29/2.93 | 5.33/4.04 |
| RXRA | 8.88/5.49 | 1.23/1.05 | 7.40/4.75 |
| TGFR1 | 7.19/5.48 | 4.20/3.27 | 6.62/5.16 |
| THRB | 4.27/3.88 | 2.99/2.66 | 4.09/3.65 |
| TRY1 | 1.58/1.41 | 3.45/3.14 | 3.43/3.18 |
| TRYB1 | 1.98/1.99 | 2.22/2.05 | 2.22/1.99 |
| VGFR2 | 1.26/1.03 | 2.93/2.43 | 2.58/2.26 |
| **AVERAGE** | **3.43/2.74** | **2.18/1.98** | **3.14/2.58** |
| **MEDIAN** | **2.77/2.65** | **2.16/2.05** | **2.58/2.26** |

**Docking Score-Based** ranking uses the lower value of the SD and ID scores.

**Table S2. Virtual screening performance of ID-rigid-AF3 evaluated at 1% cutoff.**

| System | Enrichment Factor | | # Active Hits (SD/ID/Shared) | Overlap Rate | Consensus Ranking | |
|---|---|---|---|---|---|---|
| | SD | ID-rigid | | | Docking Score-Based | Borda Count-Based |
| ACES | 3.31 | 9.49 | 15/43/3 | 4.1% | 9.49 | 9.05 |
| AKT2 | 1.69 | 0.84 | 2/1/0 | 1.4% | 0.84 | 0.84 |
| AMPC | 0.00 | 4.78 | 0/3/0 | 6.7% | 4.78 | 0.00 |
| BACE1 | 4.51 | 23.39 | 22/114/6 | 3.7% | 23.80 | 12.31 |
| BRAF | 15.46 | 12.69 | 39/32/9 | 9.6% | 12.29 | 19.43 |
| CDK2 | 8.62 | 4.75 | 69/38/21 | 11.6% | 4.87 | 12.50 |
| DPP4 | 0.74 | 1.20 | 8/13/2 | 19.1% | 1.20 | 0.74 |
| EGFR | 3.36 | 4.56 | 28/38/6 | 3.0% | 4.44 | 3.24 |
| FA10 | 8.17 | 21.88 | 65/174/52 | 26.3% | 22.00 | 16.60 |
| FABP4 | 18.73 | 25.54 | 11/15/8 | 26.7% | 22.14 | 37.46 |
| GRIA2 | 3.02 | 2.35 | 9/7/0 | 7.3% | 2.01 | 5.37 |
| HSP90A | 0.00 | 0.00 | 0/0/0 | 0.0% | 0.00 | 0.00 |
| IGF1R | 3.08 | 3.08 | 7/7/0 | 2.1% | 3.08 | 8.35 |
| ITAL | 0.43 | 17.45 | 1/41/1 | 3.3% | 17.45 | 5.53 |
| MAPK2 | 6.74 | 8.19 | 14/17/4 | 15.4% | 8.19 | 11.56 |
| MET | 1.64 | 1.23 | 4/3/0 | 0.0% | 1.23 | 4.50 |
| MK10 | 1.59 | 10.07 | 3/19/0 | 7.1% | 10.07 | 5.30 |
| MK14 | 1.64 | 15.50 | 15/142/3 | 2.7% | 15.28 | 7.42 |
| PPARD | 2.42 | 12.08 | 7/35/3 | 5.1% | 12.08 | 10.01 |
| PPARG | 5.12 | 15.21 | 37/110/25 | 11.3% | 15.35 | 14.10 |
| PTN1 | 7.07 | 6.19 | 16/14/0 | 5.2% | 6.19 | 9.28 |
| RXRA | 28.90 | 28.90 | 47/47/24 | 30.4% | 29.51 | 38.74 |
| TGFR1 | 11.69 | 2.83 | 33/8/2 | 7.8% | 3.90 | 9.56 |
| THRB | 3.60 | 7.66 | 31/66/5 | 9.2% | 7.54 | 6.96 |
| TRY1 | 2.24 | 5.13 | 17/39/2 | 6.0% | 5.13 | 2.50 |
| TRYB1 | 1.17 | 4.67 | 2/8/0 | 0.0% | 4.67 | 1.75 |
| VGFR2 | 2.73 | 7.23 | 17/45/7 | 3.1% | 7.07 | 3.70 |
| **AVERAGE** | **5.74** | **9.51** | -- | 8.5% | **9.43** | **9.51** |
| **MEDIAN** | **3.08** | **7.23** | -- | 6.0% | **7.07** | **7.42** |

**Table S3. Virtual screening performance of ID-AF3 evaluated at 1% cutoff.**

| System | Enrichment Factor | | # Active Hits (SD/ID/Shared) | Overlap Rate | Consensus Ranking | |
|---|---|---|---|---|---|---|
| | SD | ID | | | Docking Score-Based | Borda Count-Based |
| ACES | 3.31 | 6.18 | 15/28/2 | 9.4% | 7.06 | 8.38 |
| AKT2 | 1.69 | 0.00 | 2/0/0 | 2.8% | 0.00 | 1.69 |
| AMPC | 0.00 | 4.78 | 0/3/0 | 23.3% | 1.59 | 0.00 |
| BACE1 | 4.51 | 11.08 | 22/54/11 | 16.0% | 9.23 | 7.39 |
| BRAF | 15.46 | 12.29 | 39/31/8 | 13.5% | 17.05 | 15.46 |
| CDK2 | 8.62 | 6.87 | 69/55/31 | 18.5% | 6.25 | 10.87 |
| DPP4 | 0.74 | 0.56 | 8/6/2 | 27.8% | 0.28 | 0.46 |
| EGFR | 3.36 | 1.32 | 28/11/6 | 9.1% | 3.36 | 3.12 |
| FA10 | 8.17 | 16.97 | 65/135/46 | 26.8% | 15.09 | 13.96 |
| FABP4 | 18.73 | 10.22 | 11/6/5 | 26.7% | 10.22 | 25.54 |
| GRIA2 | 3.02 | 3.69 | 9/11/1 | 12.9% | 3.69 | 4.36 |
| HSP90A | 0.00 | 0.00 | 0/0/0 | 7.8% | 0.00 | 0.00 |
| IGF1R | 3.08 | 5.27 | 7/12/0 | 5.2% | 5.27 | 7.91 |
| ITAL | 0.43 | 13.62 | 1/32/1 | 4.4% | 13.62 | 2.13 |
| MAPK2 | 6.74 | 11.56 | 14/24/4 | 16.9% | 12.04 | 13.01 |
| MET | 1.64 | 1.64 | 4/4/0 | 3.4% | 2.05 | 7.36 |
| MK10 | 1.59 | 3.18 | 3/6/0 | 10.0% | 2.65 | 4.24 |
| MK14 | 1.64 | 12.44 | 15/114/1 | 5.3% | 12.11 | 5.46 |
| PPARD | 2.42 | 2.76 | 7/8/0 | 5.9% | 2.76 | 4.83 |
| PPARG | 5.12 | 7.33 | 37/53/17 | 12.4% | 7.74 | 9.40 |
| PTN1 | 7.07 | 6.19 | 16/14/0 | 7.8% | 6.19 | 0.88 |
| RXRA | 28.90 | 32.59 | 47/53/31 | 39.2% | 31.36 | 37.51 |
| TGFR1 | 11.69 | 6.73 | 33/19/6 | 18.9% | 6.73 | 13.81 |
| THRB | 3.60 | 2.32 | 31/20/1 | 25.2% | 2.55 | 1.97 |
| TRY1 | 2.24 | 1.45 | 17/11/2 | 12.5% | 1.71 | 0.92 |
| TRYB1 | 1.17 | 2.92 | 2/5/0 | 12.7% | 2.33 | 2.33 |
| VGFR2 | 2.73 | 7.71 | 17/48/6 | 3.8% | 7.39 | 3.86 |
| **AVERAGE** | **5.74** | **7.10** | -- | 14.0% | **7.05** | **7.66** |
| **MEDIAN** | **3.08** | **6.18** | -- | 12.5% | **6.19** | **4.83** |

**Table S4. Virtual screening performance of ID-NP utilizing MOE.**

| System | SD | ID | Dock Score-based Consensus Ranking |
|---|---|---|---|
| ACES | 5.52/3.18/2.47 | 4.19/2.52/2.14 | 4.85/3.27/2.43 |
| AKT2 | 0.84/1.88/2.13 | 2.53/1.88/1.88 | 2.53/1.54/2.56 |
| AMPC | 0.00/0.00/0.00 | 0.00/0.00/0.16 | 0.00/0.00/0.00 |
| BACE1 | 9.44/4.78/3.40 | 6.56/3.83/2.82 | 10.05/4.82/3.36 |
| BRAF | 2.38/1.99/1.63 | 9.51/4.86/3.51 | 7.93/4.06/2.79 |
| CDK2 | 1.50/1.28/1.10 | 1.62/1.20/0.99 | 1.75/1.23/0.99 |
| DPP4 | 0.09/0.69/0.89 | 1.11/0.89/0.89 | 1.11/0.74/0.88 |
| EGFR | 8.89/3.97/2.88 | 3.96/2.04/1.60 | 8.65/3.99/2.91 |
| FA10 | 12.20/7.32/5.28 | 4.40/3.41/2.70 | 12.07/7.07/5.23 |
| FABP4 | 0.00/0.35/0.52 | 0.00/0.35/0.35 | 0.00/0.35/0.35 |
| GRIA2 | 1.34/1.28/1.25 | 2.35/1.82/1.35 | 2.35/1.82/1.31 |
| HSP90A | 0.00/0.64/0.64 | 2.38/1.12/0.80 | 0.00/0.64/0.64 |
| IGF1R | 6.15/3.89/3.14 | 4.83/3.10/2.65 | 7.03/3.89/3.10 |
| ITAL | 1.28/0.69/0.94 | 0.43/1.63/1.67 | 0.85/0.69/0.99 |
| MAPK2 | 0.48/0.78/1.07 | 0.96/0.68/0.87 | 0.96/0.87/1.02 |
| MET | 7.36/5.24/4.06 | 4.91/2.95/2.50 | 7.36/5.24/4.02 |
| MK10 | 1.06/1.18/1.18 | 2.12/1.18/1.18 | 2.12/1.29/1.13 |
| MK14 | 3.49/2.23/1.87 | 2.62/1.62/1.48 | 3.60/2.25/1.80 |
| PPARD | 4.49/2.22/1.80 | 1.04/0.83/0.73 | 1.04/0.90/0.83 |
| PPARG | 2.90/2.35/1.80 | 2.77/2.38/1.95 | 2.90/2.16/2.00 |
| PTN1 | 19.45/7.55/4.40 | 1.77/2.31/2.00 | 13.70/7.20/4.49 |
| RXRA | 4.30/1.97/1.91 | 1.23/1.48/1.05 | 4.30/1.97/1.98 |
| TGFR1 | 1.06/2.06/2.45 | 0.71/1.28/1.85 | 1.06/1.85/2.35 |
| THRB | 6.73/4.53/3.48 | 4.76/3.44/2.95 | 5.57/4.39/3.46 |
| TRY1 | 6.84/4.67/3.64 | 5.26/3.67/3.17 | 5.26/4.17/3.44 |
| TRYB1 | 2.33/2.10/1.64 | 1.75/1.87/1.29 | 1.75/1.52/1.52 |
| VGFR2 | 3.70/3.19/2.84 | 4.98/3.03/2.66 | 3.70/3.16/2.81 |
| **AVERAGE** | **4.22/2.67/2.16** | **2.92/2.05/1.75** | **4.17/2.63/2.16** |
| **MEDIAN** | **2.90/2.10/1.87** | **2.38/1.87/1.67** | **2.90/1.97/2.00** |

**3** numbers in each cell represent the enrichment factors at **1%**, **5%** and **10%** cutoffs, respectively.
